# Striatal dopamine explains novelty-induced behavioral dynamics and individual variability in threat prediction

**DOI:** 10.1101/2021.12.21.473723

**Authors:** Korleki Akiti, Iku Tsutsui-Kimura, Yudi Xie, Alexander Mathis, Jeffrey Markowitz, Rockwell Anyoha, Sandeep Robert Datta, Mackenzie Weygandt Mathis, Naoshige Uchida, Mitsuko Watabe-Uchida

**Affiliations:** Department of Molecular and Cellular Biology, Center for Brain Science, Harvard University, Cambridge, MA 02138, USA; The Rowland Institute at Harvard, Harvard University, Cambridge, MA, USA; Department of Neurobiology, Harvard Medical School, Boston, MA, USA

## Abstract

Animals exhibit diverse behavioral responses, such as exploration and avoidance, to novel cues in the environment. However, it remains unclear how dopamine neuron-related novelty responses influence behavior. Here, we characterized dynamics of novelty exploration using multi-point tracking (DeepLabCut) and behavioral segmentation (MoSeq). Novelty elicits a characteristic sequence of behavior, starting with investigatory approach and culminating in object engagement or avoidance. Dopamine in the tail of striatum (TS) suppresses engagement, and dopamine responses were predictive of individual variability in behavior. Behavioral dynamics and individual variability were explained by a novel reinforcement learning (RL) model of threat prediction, in which behavior arises from a novelty-induced initial threat prediction (akin to “shaping bonus”), and a threat prediction that is learned through dopamine-mediated threat prediction errors. These results uncover an algorithmic similarity between reward- and threat-related dopamine sub-systems.

**Highlights:**

- Novelty-induced behaviors are analyzed using modern machine-learning methods
- Novelty induces risk assessment which develops into engagement or avoidance
- Dopamine in the tail of striatum correlates with individual behavioral variability
- Reinforcement learning with shaping bonus and uncertainty explains the data

## INTRODUCTION

Novelty induces a variety of behaviors. In the natural world, animals continuously face the problem of deciding whether to approach, avoid, or ignore a novel stimulus. Maladaptation to novelty has been implicated in anxiety, autism and schizophrenia (Baron-Cohen et al., 2005; Hirshfeld-Becker et al., 2014; Jiujias et al., 2017; Kagan et al., 1984; Orefice et al., 2016).

Behavioral responses to novelty have been modeled in different ways across fields. Within the field of reinforcement learning, novelty is often thought of as either a rewarding outcome or a predictor of a potential reward, thereby prompting exploration before the first rewards are received (Kakade and Dayan, 2002; Xu et al., 2021). In this way, it can be incorporated into existing reinforcement learning frameworks. Similarly, artificial intelligence models have been created that are “curious” or intrinsically motivated (Colas et al., 2019; Oudeyer et al., 2007, 2016; Stout et al., 2005). Some of these models use information gain, a reduction in the difference between the current event and what was expected over time, to define event novelty (Jaegle et al., 2019; Kaplan and Oudeyer, 2007). Notably, while many computational models of novelty capture the neophilic aspects of novelty behavior, they fail to capture the neophobia and the interplay between approach and avoidance in response to novelty, observed in natural novelty responses.

Dopamine has long been thought to be a critical regulator of reward-oriented behaviors, and electrophysiology studies have shown that dopamine signals the discrepancy between actual and predicted reward value (Montague et al., 1996; Schultz et al., 1997). In reinforcement learning, dopamine can be used as an evaluation signal to reinforce a rewarding action (reinforcement learning). However, recent studies have found that some dopamine neurons are activated by novelty (Horvitz et al., 1997; Lak et al., 2016; Ljungberg et al., 1992; Menegas et al., 2017, 2018; Morrens et al., 2020; Schultz, 1998). To incorporate these novelty responses into the reinforcement learning framework, it has been proposed that dopamine novelty response may correspond to optimism or the potential for reward (Kakade and Dayan, 2002).

Although it has been widely assumed that dopamine neurons broadcast reward prediction error signals to a wide swath of targets, recent studies have shown that dopamine neurons projecting to different targets send distinct information (Kim et al., 2015; Lerner et al., 2015; Menegas et al., 2017; Parker et al., 2016). Importantly, the canonical dopamine system – those that project from the ventral tegmental area (VTA) to the ventral striatum (VS) – does not respond to novel stimuli at the population level (Menegas et al., 2017). Recent studies in monkeys also found that dopamine neurons in substantia nigra pars compacta (SNc) do not respond to novelty per se (Ogasawara et al., 2021), but rather respond to novelty in the context of information seeking for reward (Bromberg-Martin and Hikosaka, 2009). In contrast, recent studies found that dopamine neurons that project to the tail of the striatum (TS) or the prefrontal cortex play a role in task-independent novelty-related behaviors (Menegas et al., 2018; Morrens et al., 2020).

A recent study found that dopamine in the posterior tail of the striatum (TS) displays unique response properties (Kim and Hikosaka, 2013; Menegas et al., 2017). TS-projecting dopamine neurons are strongly activated by high intensity or novel external stimuli in the environment (Menegas et al., 2017, 2018), or by salient visual cues, but not by reward (Kim et al., 2015).

Functionally, TS-projecting dopamine neurons facilitate avoidance of a threatening stimulus including a novel object (Menegas et al., 2018).

However, how dopamine modulates novelty-driven behaviors is not clearly understood, as there are several limitations in previous studies. Firstly, previous studies treated novelty-related behavior as a binary choice of either approach (orient, saccade) or avoidance, and often ignored the behavioral complexity, dynamics and individual variability, which is essential to understand the computations underlying novelty responses. Variability in the novelty-triggered behavioral data had been even interpreted as experimental deficits (Corey, 1978). However, individual variability is an important factor to understand the neural computations (Marder and Goaillard, 2006). Secondly, many previous studies were conducted in constrained environments that limited behavioral choices (Menegas et al., 2017; Morrens et al., 2020; Ogasawara et al., 2021). It has been reported that animals respond differently to novel objects depending on whether the animal is in a small environment (“forced exposure”) or in a sufficiently large enclosure to be able to choose between exploring or totally avoiding a novel object (“voluntary exploration”) (Corey, 1978; Rebec et al., 1997). Thirdly, the definition of novelty varies across studies. Recent studies emphasized the computational difference between stimulus novelty and contextual novelty: the former refers to the quality of not being previously experienced or encountered, and the latter refers to the “surprise” when what is experienced does not match with what was expected in time and/or context (i.e. prediction error) (Barto et al., 2013; Kumaran and Maguire, 2007; Ranganath and Rainer, 2003; Xu et al., 2021). Finally, novelty avoidance or neophobia is typically characterized as mal-adaptative behavior but not a sensible action.

In this study, we used machine learning to characterize individual variability in behavioral novelty responses while mice freely explored a novel object placed in a large arena. We subsequently examined the effects of two types of novelty, the first in which a mouse explored a new stimulus (“stimulus novelty”) and the second in which a mouse explored a familiar stimulus in a new location (“contextual novelty”). These different novelty manipulations induced distinct patterns of behavior, which were differentially affected by ablation of TS-projecting dopamine neurons. The diversity and the dynamics of the observed novelty behaviors were well captured by a simple reinforcement learning model, which incorporates the concepts of initial estimation (“shaping bonus”) and uncertainty. We propose that novelty avoidance is a critical defensive strategy in which a novel stimulus causes default estimation of potential threat when the outcome is unknown. Because death or significant injury prevent learning, the brain may have adapted to estimate the degree of threat posed by a novel object through its physical salience, signaled by dopamine in TS.

## RESULTS

### Diverse behavioral responses to novelty

To better study the diversity of novelty-related behavior, we designed an open arena novelty exploration task (Figure 1). A novel object was placed in the corner of a large (60cm-by-60cm) arena and mice were allowed to move freely and interact with the object at will (Figure 1a-c). Mouse movements were captured using an overhead camera that recorded four channels: three color channels (RGB) and one channel for depth (Xbox Kinect; see Methods). We used DeepLabCut (Mathis et al., 2018) to track the nose, ears, and tail base of the mouse (see Methods). This tool provides access not just to basic center-of-mass tracking (used commonly in previous studies; Menegas et al., 2018) but multi-point measurements, a benefit that will be leveraged in our analysis.

**Figure 1.**
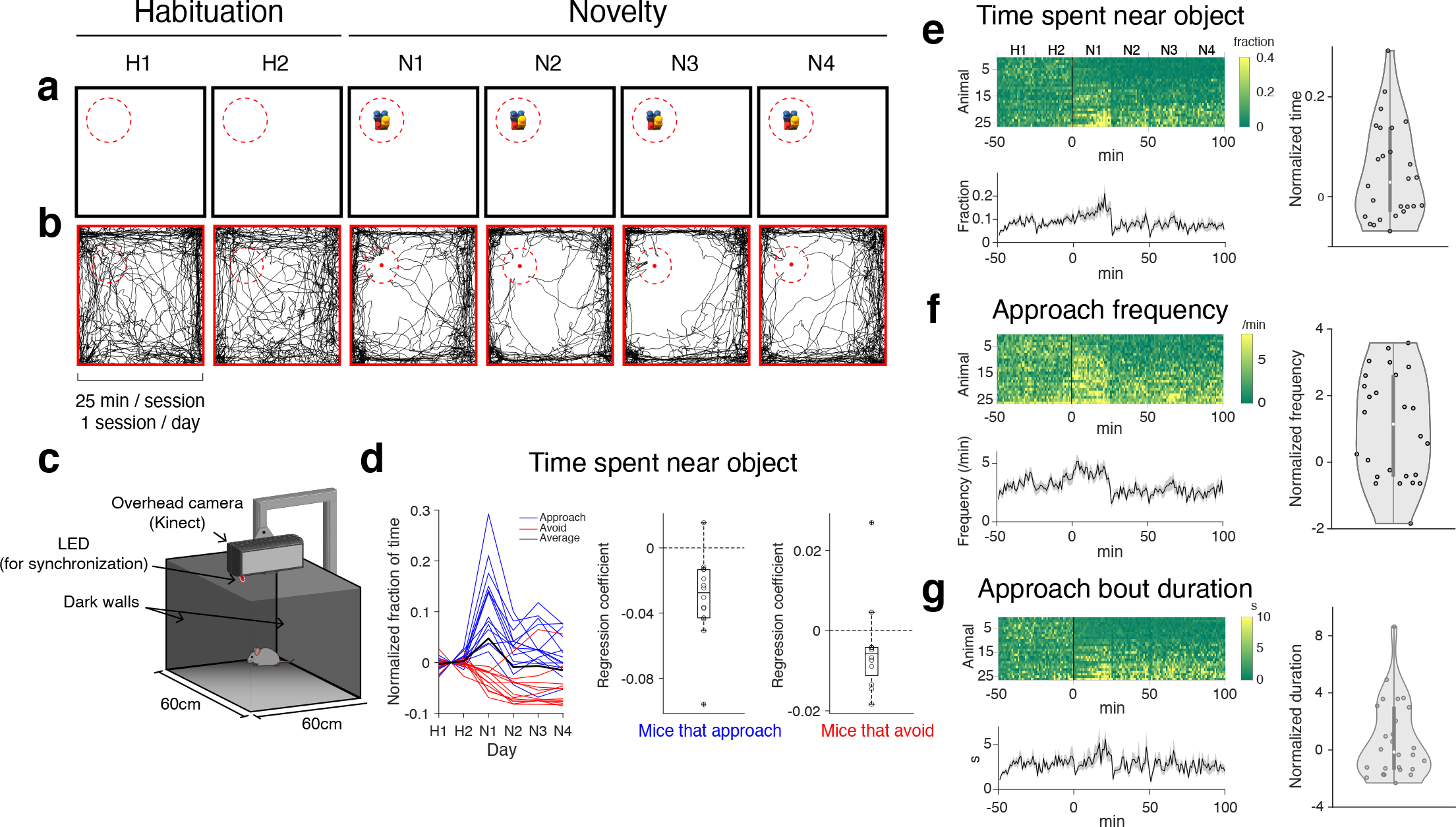
Diversity of novelty behavior is captured in open arena **a.** Experimental paradigm for novelty exploration. Mice are habituated to arena with no object present (days H1/H2) before testing response with a novel object present (days N1-N4). Mice have never encountered this object before. Dotted red lines indicate area of analysis. **b.** Representative trajectory of nose position from a single animal in the first 10 minutes of each session. Dotted red lines show area of analysis (7cm radius). **c.** Arena setup, consisting of 60cm-by-60cm open arena with dark walls (darkness was important for video recording for MoSeq), an overhead camera (Xbox Kinect for Windows; captures rgb and depth video), and an LED light for photometry synchronization. **d.** Left: Time spent within object area (7cm radius; subtracted with average time during habituation H1/H2) across experimental days. Colors denote whether mice approached (above 0) or avoided (below 0) the object area on the first day of novelty (N1). Average value across mice is shown in black. Middle/right: Regression coefficients of time spent near object with novelty days for each animal (N1-N4; p = 6.1*×*10^-4^, n=14, approach mice; p = 0.19, n=11, avoidance mice, t-test compared to 0). **e.** Left: Fraction of time spent within object area (radius=7cm) over 6 days. Zero on the x-axis indicates the beginning of the first day of novelty, and habituation sessions are indicated by negative values on the x axis. Bottom plot shows mean *±* SEM (n=25). Right: Fraction of time near object on first day of novelty for each mouse (subtracted with average on days H1/H2). **f.** Frequency of approaches across sessions. Same format as in **e**. **g.** Duration of approach bouts across sessions. Same format as in **e**.

Novelty sessions consisted of 25 minutes of exploration per day over the course of several days. Prior to testing, mice were habituated to the empty arena for two days. When no object was present, mice explored all parts of the arena (Figure 1b). Once a novel object was introduced, mice changed their behavior (Figure 1b, N1-N4), exhibiting many bouts of approach (defined as epochs in which the mouse was in close proximity to the object, see Methods) followed by retreat.

On the first day of novelty (N1) when the mice were first encountering the object, mice either spent more time (approach) or less time (avoid) within the object area compared to habituation days (Figure 1d, left). Mice that spent less time than previous days near the object on N1 (Figure 1d, red traces, 12/26 animals) tended to continue avoidance on subsequent days (11/12 animals, Pearson’s correlation coefficient R=0.70, p=6.3*×*10^-5^, N1 vs N2, n=26 animals), while the increase in time spent near object on N1 by some mice (Figure 1d, blue traces, 14/26 animals) tended to be transient compared to subsequent days. We quantified this trend by performing a linear regression of time spent near the object with novelty days (N1-N4). Mice that favored approach (Figure 1d, middle) showed significant decreases in time spent near the object across days (linear regression of time spent near object with day, beta coefficient vs 0, p=6.1*×*10^-4^, t-test). This finding suggests that for this subset of mice, novelty decreases over time. Thus, mice exhibit variable responses to novelty that is obscured by averaging (Figure 1d, black line).

Time spent near object is a function both of the frequency with which mice approach the object (approach frequency, Figure 1f) and how long they spend near the object once they approach (approach bout duration, Figure 1g). Both approach frequency and approach bout duration were variable across animals, with prominent similarity in the dynamics between approach bout duration and time spent near object (Figure 1e-g); Similar to time spent near object, some mice showed longer approach bout duration than previous days on N1, while some mice showed shorter bout duration across all entire sessions (Figure 1g).

Together, we observe a diversity in novelty responses: some mice react to the novel object by approaching, and some by avoiding. The results demonstrate that “approach” mice show a decrease in approach over time (closer to pre-novelty baseline), suggesting that approach behavior is a response to novelty, whereas avoidance tends to be sustained.

### Novelty triggers stereotypical risk assessment response

Close examination of the trajectory of nose and tail while a mouse is within the object area revealed that in the first several approach bouts, mice oriented themselves to face the object (Figure 2). While their time spent near the object fluctuated every day (Figure 1), this orientation behavior was not seen at the beginning of subsequent novelty days (Figure 2a, N2-N4). As a result, when the animal reached the closest point to the object, the closest body part was always the nose, not the tail (Figure 2c, N1), which was also unique in the first approach bouts (first ∼50 bouts in an example animal). These data suggest that the novelty response characterized by “approach with tail behind” is unique to early interactions with a novel object.

**Figure 2.**
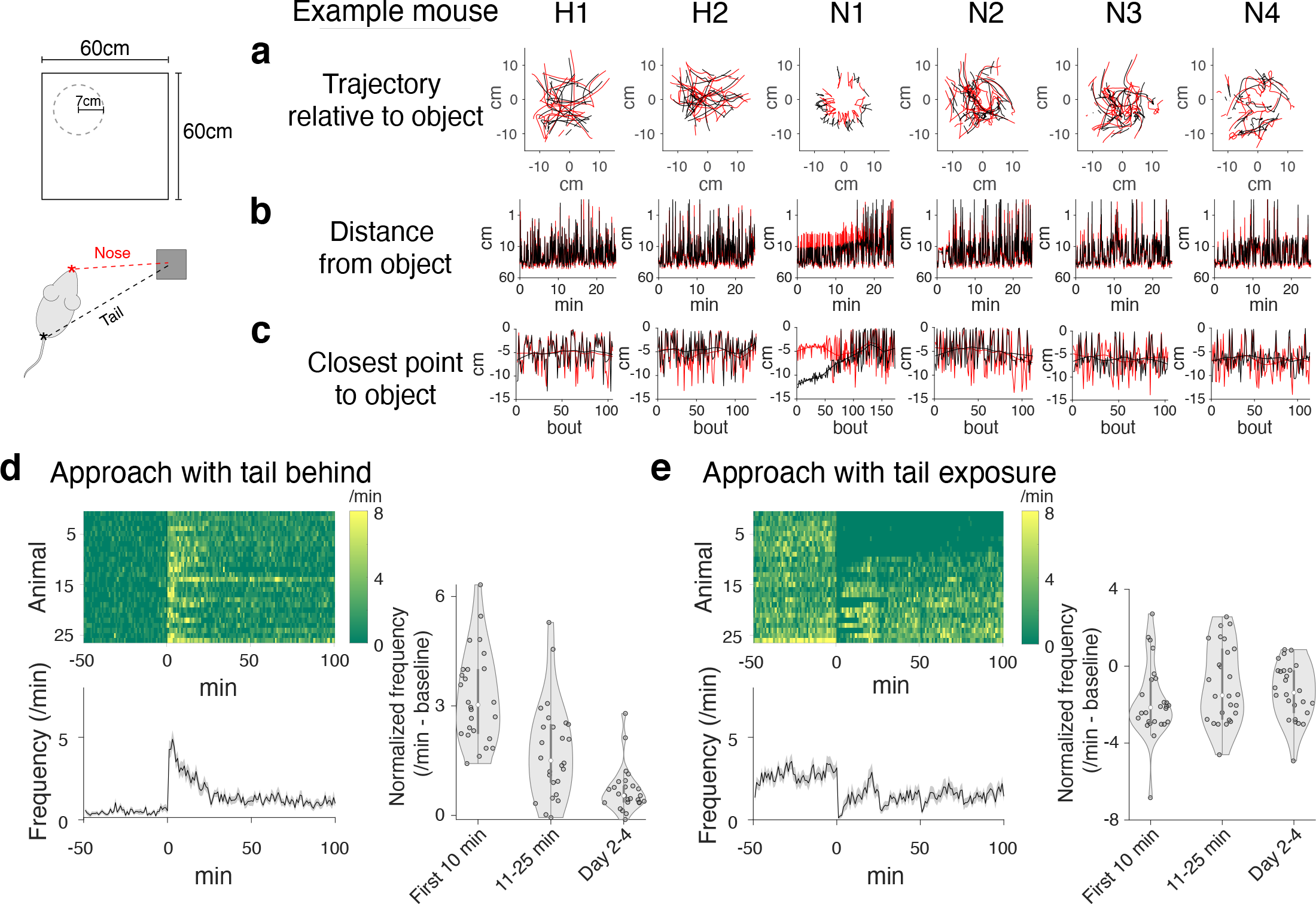
Stereotypic behavioral response to novelty. **a.** Representative trajectory of position of nose or tail (in red and black, respectively) from a single mouse. First 20 bouts of each session are shown. **b.** Nose and tail position relative to object in a single animal. **c.** The closest position to object within each bout for nose and tail in a single animal. **d.** Left: Frequency of approach bout with tail kept behind nose (per minute) over 6 days. Black line (zero on the x-axis) indicates the beginning of the first day of novelty, and habituation sessions are indicated by negative values on the x axis. Bottom plot shows mean *±* SEM. Right: Breakdown of average frequency of approach with tail behind (normalized with baseline on habituation) for each mouse in the beginning of N1 (left), the end of N1 (middle), and for novelty days 2-4 (right). Tail behind approach frequency decreases over time (p=2.8*×*10^-11^, t-test, n=26, beta coefficients of linear regression of frequency vs time). **e.** Left: Fraction of tail exposure across sessions. Right: Breakdown of average frequency of approach with tail exposure (habituation normalized). Same format as in **d**. No significant linear change in tail exposure across time (p=0.20, t-test, n=26, beta coefficients of linear regression of frequency vs time).

To quantify this prominent novelty-related behavior we classified approach bouts based upon orientation, which revealed that every mouse approached the object with the tail behind in the first 10 min on the first day of novelty (Figure 2d, n=26). The frequency of approach with tail behind decreased over time (Figure 2d, linear regression of frequency with time, beta coefficient vs 0, p=2.8*×*10^-11^, t-test). Over the course of the first day, some mice started to allow their tails to be exposed to the object, while some mice did not expose their tails to the object during entire sessions (Figure 2e). Among the mice exhibiting tail exposure, some showed high frequency of tail exposure transiently in the later time on N1 (Figure 2e right, 11-25min).

These results thus revealed a robust stereotyped response at the beginning of interactions with a novel object, one that resembles a form of behavior described as “risk assessment” (Blanchard et al., 1991; Gottlieb and Oudeyer, 2018; Kidd and Hayden, 2015). On the other hand, post-assessment behaviors were diverse, with individual animals exhibiting a wide spectrum of approach or avoidance behaviors, with approach gradually decreasing over days. We operationally refer to post-assessment approach as “tail exposure engagement,” to distinguish from risk assessment.

### Post-assessment engagement is suppressed by stimulus novelty

Initial encounters with a novel object inevitably include both stimulus novelty and contextual novelty. First, an animal might react by identifying the object as one it has never experienced in its life. In this case, the brain has to search its stored memory of all objects in the past to detect novelty (stimulus novelty) (Barto et al., 2013). In addition to stimulus novelty, an animal might be surprised because the object was not expected in the current context (Barto et al., 2013). In this case, the brain has to compare the current state with the predicted state to detect novelty (contextual novelty) (Ranganath and Rainer, 2003).

To separate the impact of different kinds of novelty on behavior, we incorporated object pre-exposure into our behavioral paradigm (Figure 3), which dramatically changed the animal’s reaction to the object in test sessions. As shown above (Figure 1, Figure 2), some mice spend more time near a novel object on N1, while others spend less (Figure 3b, left). In contrast, mice consistently approached an unexpected familiar object (Figure 3b, middle). As a population, mice with an unexpected familiar object spent significantly more time near the object than mice with a novel object (p=0.018, Kolmogorov-Smirnov [K-S] test), and also exhibited limited tail behind approach and quickly switched to tail exposure (Figure 3d). Mice interacting with a novel object used tail-behind approach significantly more frequently than mice with an unexpected familiar object (Figure 3c-e, p=0.0031, t-test) and used tail exposure approach significantly less frequently (Figure 3c-e, p=0.0031, t-test).

**Figure 3.**
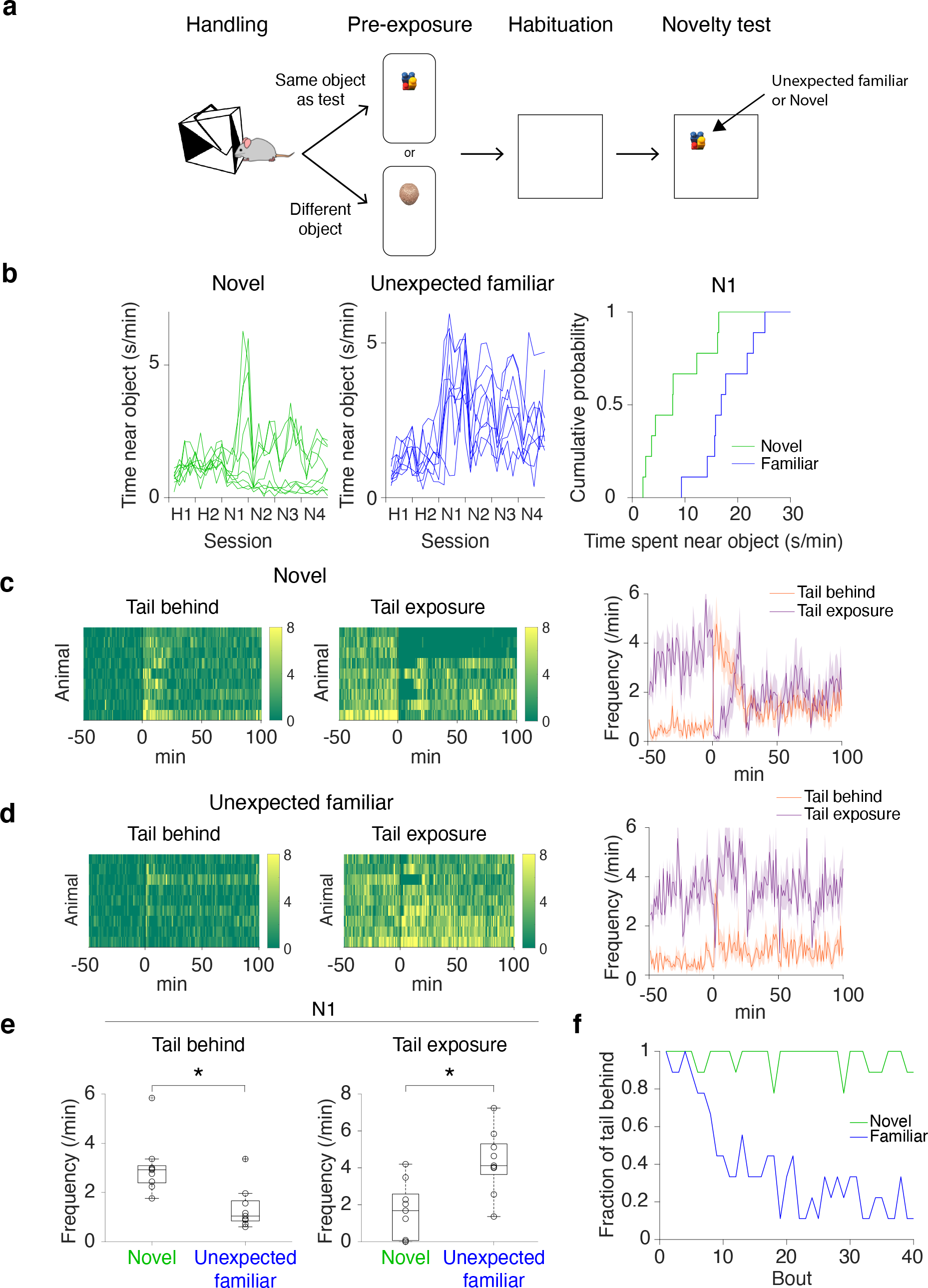
Suppression of post-assessment engagement with stimulus novelty. **a.** Experimental setup. Handling, habituation, and testing are kept constant between the two novelty groups. Pre-exposure differs: one group of mice are pre-exposed to the test object (unexpected familiar) and another group of mice to an object different from the test object (novel). **b.** Left/middle: Frequency of time spent near a novel (left, green) or unexpected familiar (middle, blue) object across sessions. Right: Cumulative probability of each group of mice spending certain amounts of time near object on the first day of novelty (N1). Mice spend less time overall near a novel object (p=0.018, n=9 animals for each group, Kolmogorov-Smirnov test). **c-d.** Left: Frequency of approach with tail behind and approach with tail exposure towards novel (c) and unexpected familiar (d) objects. Right: Comparison of approach types (mean *±* SEM). **e.** Frequency of approaches on the first day of novelty (average across whole session for each mouse). Approach with tail behind is more frequent towards novel objects (p=0.0031), whereas approach with tail exposure is more frequent towards unexpected familiar objects (p=0.0031, n=9 animals for each group, t-test). **f.** Fraction of animals with approach with tail behind towards novel or unexpected familiar objects in each approach bout.

Our observation that tail-behind approach was consistently observed at the beginning of N1 in both groups (Figure 3f) suggested that risk assessment behavior is driven by unexpectedness, not specifically by stimulus novelty. However, in response to an unexpected familiar object, mice exhibited a quick transition to approach with tail exposure (engagement), suggesting that stimulus novelty suppresses engagement.

### Ablation of TS-projecting dopamine neurons biases post-risk assessment behavior towards approach

To understand the computational role of dopamine in TS in novelty-driven behaviors, we performed ablation of TS-projecting dopamine neurons with 6-hydroxydopamine (6OHDA) (Figure 4, Figure S1). Consistent with our previous study (Menegas et al., 2018), animals with ablation of TS-projecting dopamine neurons spent more time near a novel object than animals with injection of a vehicle (Figure 4c, ablation vs sham, p=0.030; K-S test) and showed longer duration of approach bouts (Figure S2). When analyzing risk assessment and engagement, all sham-lesioned animals exhibited approach with tail behind at the beginning (Figure 4d-e, n=17 animals). As a population, the frequency of tail behind approach was strikingly high on N1 with limited tail exposure approach (Figure 4d, right). Ablation mice also expressed early risk assessment behaviors. In fact, all ablation mice expressed approach with tail behind in early periods of N1 (Figure 4d-e, left, n=17 animals). After risk assessment, more ablation mice showed transition to tail exposure approach, resulting in higher frequency of tail exposure as a population (Figure 4d-f, p=0.010, sham vs ablation, t-test).

**Figure 4.**
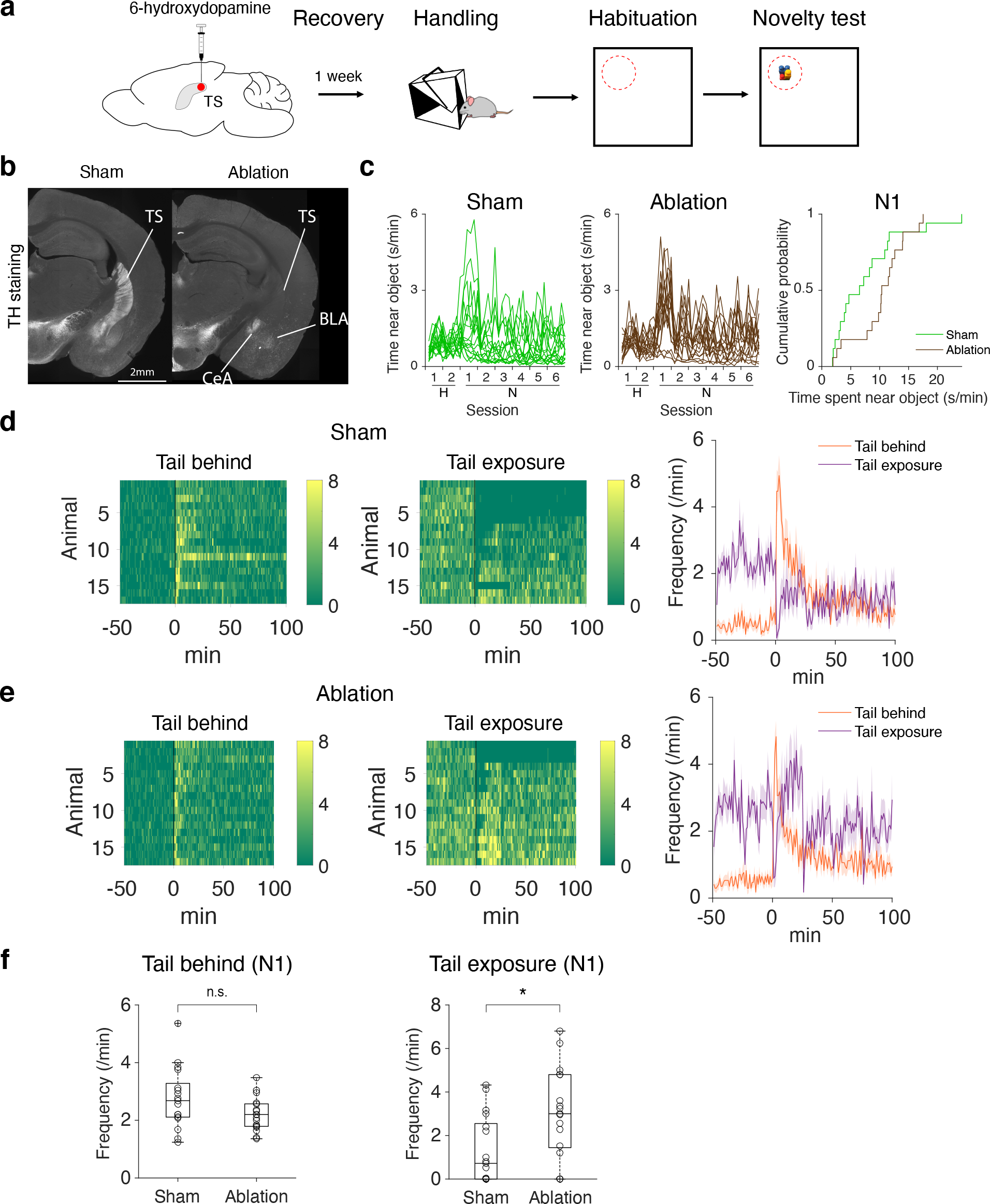
Ablation of TS-projecting dopamine neurons promotes post-assessment engagement. **a.** 6OHDA was injected bilaterally into TS. **b.** Representative images of coronal sections (bregma = -1.5) from sham (left) and ablation (right) animals, stained with anti-tyrosine hydroxylase (TH) antibody. BLA: basolateral amygdala; Ce: central amygdala. **c.** Left/middle: Frequency of time spent near object for sham (left, green) and ablation (right, brown) mice across sessions. Right: Cumulative probability of each group of mice spending certain amounts of time near novel object on the first day of novelty (N1). ablation vs sham, p=0.030 (K-S test, n=17 animals for each group). **d.** Frequency of approach bouts with tail behind or exposure for sham mice. Right: mean *±* SEM, n=17. **e.** Frequency of approach with tail behind or exposed for ablation mice. Right: mean *±* SEM, n=17. **f.** Comparison of average frequency of approach with tail behind (left; p=0.069, n=17 for each, t-test) and approach with tail exposure (right, p=0.010, n=17 for each, t-test) on N1.

These results demonstrate that ablation of TS-projecting dopamine neurons increased approach with tail exposure, suggesting that intact dopamine in TS suppresses post-assessment engagement, resulting in premature transition to engagement in ablation mice.

### Dissection of novelty-driven behaviors by behavioral segmentation

So far, we classified approach types by focusing on animal’s tail position relative to nose. To segment behavioral responses to novel objects into constituent components, we next analyzed the same data using MoSeq (Wiltschko et al., 2015), an unsupervised machine learning-based behavioral characterization method, that identifies behavioral motifs or “syllables” from depth imaging data (Figure 5). We first asked whether certain syllables were overrepresented within approach bouts, and noticed that some syllables were frequently expressed near the time of retreat. One syllable stood out (syllable 79, purple) in both the novel object mice and the sham mice. To examine whether any of the syllables were frequently and specifically expressed in different novelty conditions, or with novel or unexpected familiar objects, we first identified a set of syllables that was both highly used and which exhibited significant differential expression across groups (p<0.05, ANOVA; n=4 conditions). Among these 12 syllables, there were 3 that were enriched in the novel object condition and 4 that were enriched in the unexpected familiar object condition (Figure S3).

**Figure 5.**
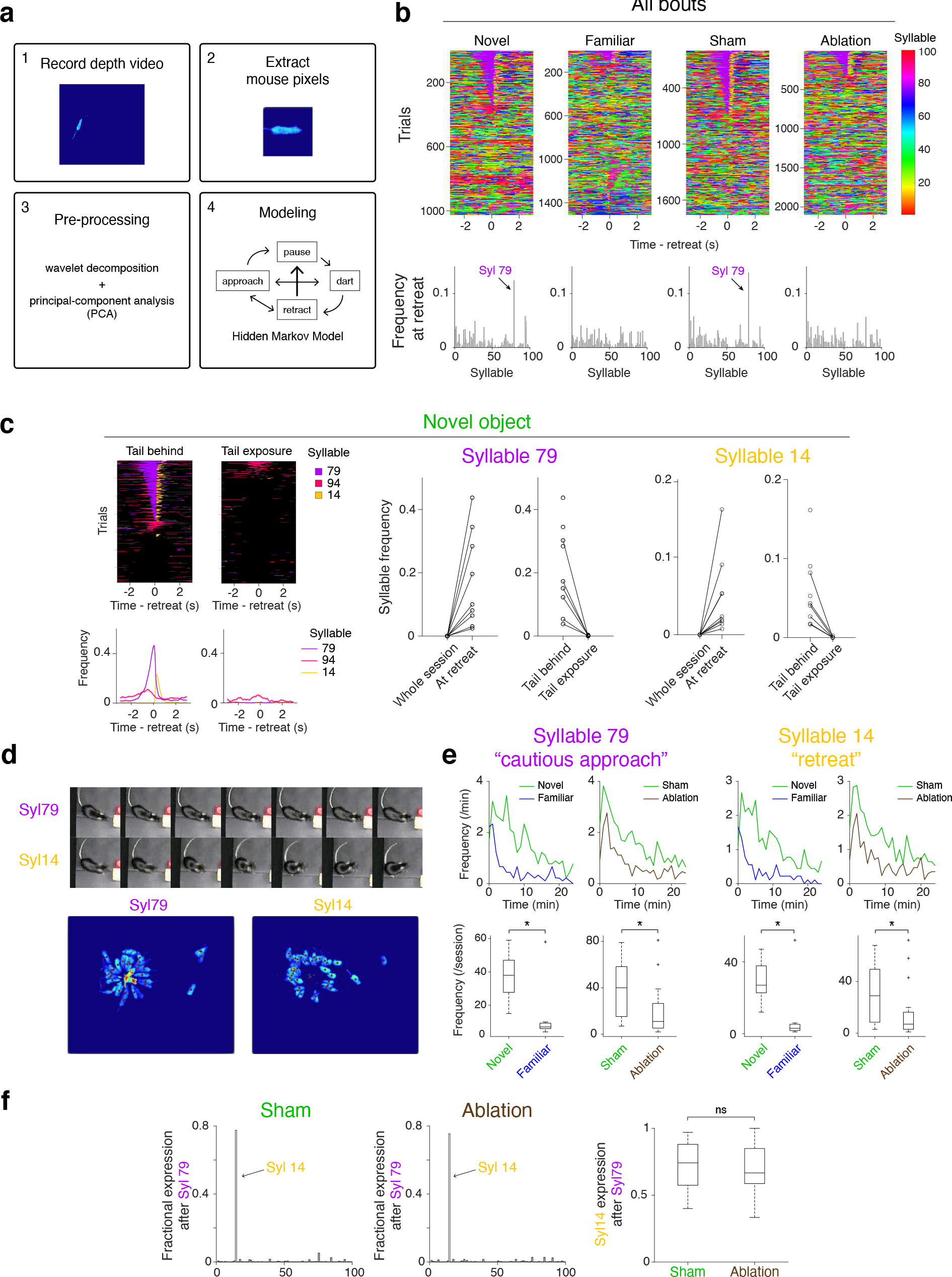
Behavioral segmentation of novelty responses using MoSeq **a.** MoSeq workflow for extracting behavioral syllables from raw depth video (see Wiltschko et al. 2015 for detailed methods). **b.** Top: Frequency of syllable usage across approach bout trials, aligned to time of retreat (n=100 syllables, output from MoSeq), on the first day of novelty (N1). Bottom: Average syllable usage at the time of retreat (-1s to 1s) for each syllable (averaged across all trials in all mice of the same condition). **c.** Left top: Expression of syllables enriched with a novel object (see Supplemental Figure 3). Colors in heatmap indicate when syllables of interest were expressed, and times when any other syllables were expressed are in black. Left bottom: Average frequency of syllable usage at the time of retreat. Middle and Right: Frequency of syllable 79 and 14 usage. **d.** Top: Image series for example cautious approach bout (syllable 79, top row) and example retreat bout (syllable 14, bottom row). Bottom: Still frames from crowd videos for syllable 79 and 14 (full videos in Supplemental Video S1 and S2). Object is in top left corner of each image. Mice are in light blue, and red dots are superimposed on image for each frame when mouse expresses the syllable of interest. Multiple instances of the syllable being expressed are superimposed on each other. **e.** Average frequency of syllable usage at any time/location on N1 for syllable 79 (left two columns) and syllable 14 (right two columns). Top: Time course of syllable expression averaged across animals within each condition, comparing either novel and familiar object mice or sham and ablation mice. Bottom: Average syllable expression on N1. (novel object vs unexpected familiar object, p=4.9*×*10^-4^, syllable 79; p=4.9*×*10^-4^, syllable 14, n=9 animals for each; sham vs ablation, p=0.010, syllable 79; p=0.030, syllable 14, n=17 animals for each, K-S test) **f.** Left: Fractional expression of each syllable after syllable 79 in sham animals. Right: Fractional expression of each syllable after syllable 79 in ablation animals. Both sham and ablation animals show prominent expression of syllable 14 after syllable 79. Right: Fraction of syllable 14 expression following syllable 79 expression in sham and ablation animals. Ablation animals show no difference in level of syllable 14 usage after syllable 79 (p=0.72, n=17 animals for each, t-test).

We next examined the relationship between the expression of these syllables and our classification of approach types: approach with tail behind and approach with tail exposure. We focused on 3 syllables that were enriched in the novel object condition; when we divided approach into the two types, we found that syllable 79 and 14 were almost exclusively expressed in approach with tail behind (Figure 5c, usage was 46.3% and 22.9% of all approach with tail behind (n=684) for syllables 79 and 14, respectively). Both syllables 79 and 14 were highly enriched at the time of retreat compared to the whole session, nearly always occurring during bouts with tail behind rather than with tail exposure (Figure 5c). Interestingly, syllable 79 was expressed just before the time of retreat and was reliably followed by syllable 14 (14 follows 79, 71.3%*±*18.9 of usages, mean*±*SEM, n=17 sham animals, Figure 5f, left).

Visual inspection of the videos (Video S1, Video S2) and video clips (Figure 5d, Figure S4) revealed that syllable 79 represented a “cautious approach” pattern and that syllable 14 represented a “cautious retreat” pattern. These results indicate that cautious approach and retreat are linked, and together make up risk assessment behavior. Thus, two out of three syllables enriched in the novel object condition were related to risk assessment behavior, which is consistent with our observations made through body part tracking (DeepLabCut) demonstrating that approach with tail behind is more pronounced with a novel object (Figure 3).

Consistent with the temporal dynamics of risk assessment characterized above, syllables 79 and 14 showed a gradual decay in usage over the course of the first day of novelty (Figure 5e). Both syllable 79 and syllable 14 were expressed more frequently over the course of the session with a novel object compared to an unexpected familiar object (novel object vs unexpected familiar object, p=4.9*×*10^-4^, syllable 79; p=4.9*×*10^-4^, syllable 14, K-S test). Interestingly, syllables 79 and 14 were also expressed more frequently in sham mice compared to ablation mice (sham vs ablation, p=0.010, syllable 79; p=0.030, syllable 14, K-S test). Thus, ablation of TS-projecting dopamine neurons decreased both novelty responses and usage of risk assessment syllables 79 and 14, although our manual classification using DeepLabCut could not detect the small difference (Figure 4f). On the other hand, syllables that were enriched in the familiar object group were not enriched in ablation mice (Figure S3).

Our finding that the expression of both syllables 79 and 14 were decreased in ablation mice indicates that TS dopamine impacts both cautious approach and retreat behaviors. This is surprising because if approach and retreat are opposing behaviors, and dopamine in TS reinforces only retreat, ablation of TS-projecting dopamine neurons should predominantly affect retreat. However, the specific syllables associated with approach and retreat were both affected by ablation. We next compared transition from syllable 79 to 14 in sham and ablation animals. Transition from syllable 79 to syllable 14 was similarly high in both animal groups (Figure 5f), indicating that choice of retreat types, characterized by a combination of syllables 79 and 14, was already determined before approach. Ablation of TS-projecting dopamine neurons decreased risk assessment, characterized by a specific posture of approach-retreat, but did not change the structure of risk assessment behaviors, characterized by the sequence of unique syllables.

### TS dopamine response to novelty reflects individual variability in behavior

To better understand the role that TS dopamine plays in novelty behavior, we monitored dopamine release in TS using fiber fluorometry (photometry) with a dopamine sensor (GRAB-DA2m, (Sun et al., 2018, 2020)) while animals explored a novel object (Figure 6). Consistent with our previous observations of dopamine axon calcium in TS (Menegas et al., 2018), we observed dopamine release in TS around the time of retreat onset (Figure 6c, Figure S5). We also examined dopamine release at the start of approach and at the end of retreat. Dopamine was specifically released when animals were at the closest point from an object (Figure 6c), consistent with the idea of risk assessment or evaluation.

**Figure 6.**
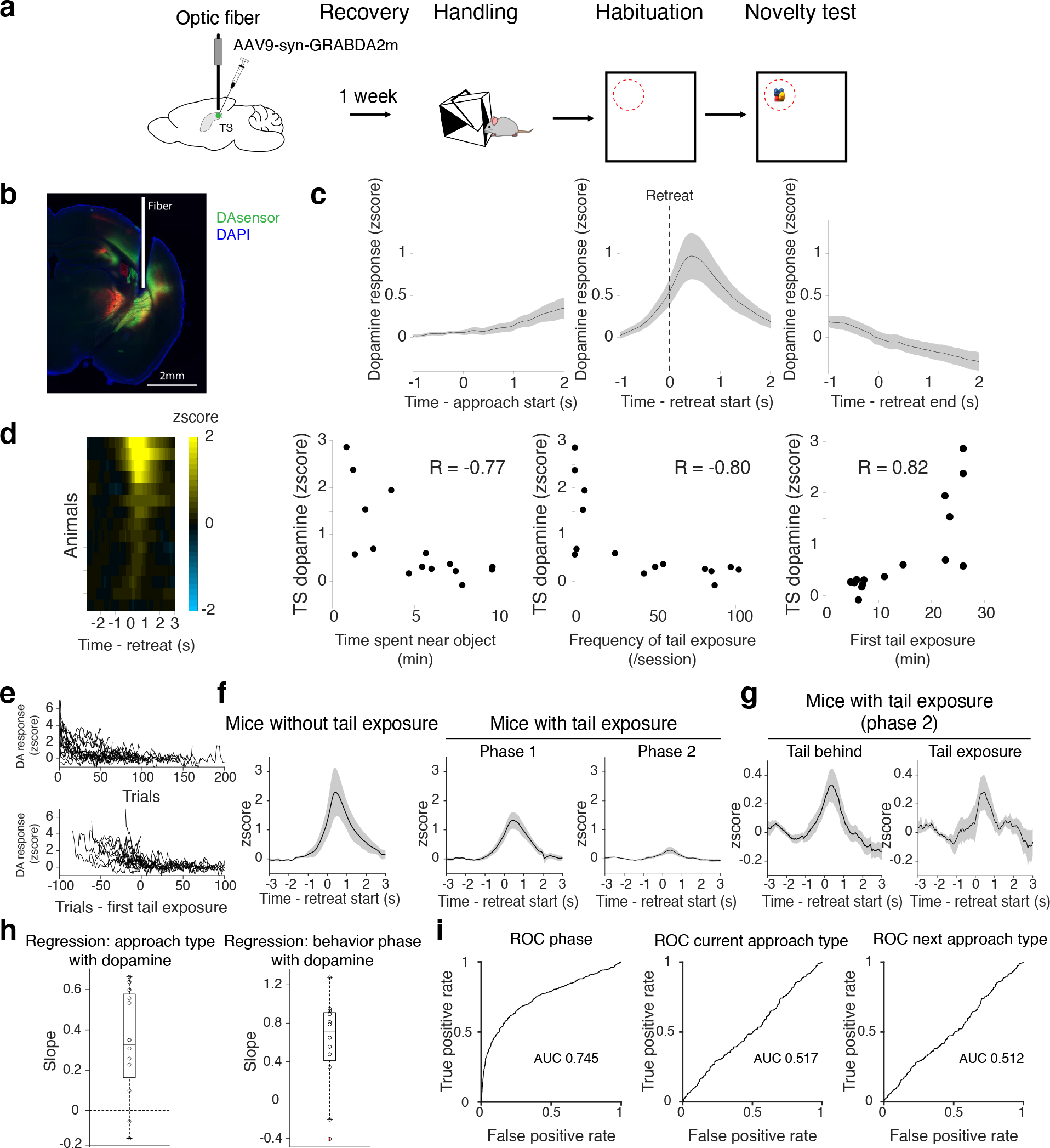
Individual variability in behavior correlates with dopamine in TS. **a.** Experimental procedure for photometry recording. AAV for dopamine sensor was injected into TS unilaterally and then an optic fiber was implanted at the same site (see Methods). **b.** Example recording site of dopamine axons (DA sensor expression in green, tdTomato in red, DAPI in blue). **c.** Dopamine signals aligned to approach start (left), retreat start (middle) and retreat end (right). mean *±* SEM, n=15. **d.** Far left: Dopamine signal on N1 aligned to time of retreat. Each row is the average signal across all retreats for a single mouse. Right: Average dopamine signal of each animal plotted against time spent near the object (second from left), frequency of tail exposure (second from right), or time of first tail exposure in session (far right). Higher dopamine signal correlates with less time spent near the object (Spearman’s correlation coefficient R=-0.77, p=0.0012, n=15 animals), more tail exposure (R=-0.80, p=2.8*×*10^-4^), and delayed tail exposure (R=0.82, p=1.7*×*10^-4^). First tail exposure for mice that never showed tail exposure (3 animals) was set to 25min, the last time point. **e.** Top: Time course of dopamine activity across trials for each animal. Bottom: Time course of dopamine activity aligned to first tail exposure. **f.** Dopamine signal in mice that never showed approach with tail exposure (left, n=3) and those that approach with tail exposure (middle/right, n=12). Trials in animals with approach with tail exposure were divided into phase 1 or phase 2 by time of first tail exposure. mean *±* SEM. **g.** Dopamine signal during phase 2 in mice that express tail exposure. Signal is aligned to retreat of tail behind approach (left) or retreat of tail exposure approach (right). Mean *±* SEM, n=12. **h.** Beta coefficients from regression of dopamine signal against approach type (left, p=0.0034) and behavioral phase (right, p=0.0034, t-test with 0, n=12). **i.** Left: Receiver operating characteristic (ROC) curve evaluating performance of classification of behavior phase with dopamine activity (AUC = 0.74). Middle/right: ROC curves evaluating classification of current approach type (middle; AUC = 0.517) or next approach type (right; AUC = 0.512) with dopamine activity.

As described above (Figure 1, Figure 2), behavioral responses to novelty were variable across animals. Interestingly, the dopamine responses observed in response to a novel object were also variable (Figure 6d, left). We tested whether the individual variability of dopamine responses corresponded to individual variability in behavior (Figure 6d). We found that mice with high average TS dopamine responses on N1 tended to spend less time near the object (Figure 6d, second from left; Pearson’s correlation coefficient R=-0.77, p=0.0012), showed less frequent tail exposure (Figure 6d, second from right; R=-0.80, p=2.8*×*10^-4^), and had slower transition to the first approach with tail exposure (Figure 6d, right; R=0.82, p=1.7*×*10^-4^).

We next examined whether dopamine responses and behaviors are correlated within each animal. We first tested correlation between dopamine responses and current approach types (tail behind or tail exposure). Dopamine responses were significantly correlated with approach types (Figure 6h, left, responses vs risk assessment, p=0.0034, t-test). However, the trial-to-trial correlation detected in this analysis may be caused by a global correlation, not necessarily by trial-based correlation, since both dopamine responses and approach types have a global trend (both dopamine responses and approach with tail behind decrease over time). We therefore examined whether the dopamine response correlated with the animal performing early risk-assessment phase (phase 1) or the late engagement phase (phase 2), which was defined based on the time when mice first expressed an approach with tail exposure (Figure 6e-f). On average, dopamine response was higher in phase 1 than in phase 2 (p=0.0059, n=12 animals, paired t-test, Figure 6f, right); note that some mice never showed approach with tail exposure (and we therefore excluded these animals from this analysis) but that they generally exhibited strong response during approach without tail exposure (Figure 6f, left). Linear regression revealed a significantly positive correlation between dopamine responses and behavior phases (Figure 6h, right, responses vs phase 1, p=0.0034). Furthermore, receiver operating characteristic (ROC) analysis revealed that dopamine responses in TS predicted the behavior phase with high accuracy (Figure 6i, left, area under curve (AUC)=0.745). We next examined the relationship between dopamine responses and approach types on a trial-by-trial basis within phase 2 (because phase 1 includes only one approach type). On average, dopamine responses were similar between approach types in phase 2 (p=0.90, n=12 animals, paired t-test Figure 6g). We performed ROC analysis and found that a dopamine response does not effectively predict a current or the next approach type (AUC = 0.517 and 0.512, Figure 6i, middle and right). These results indicate that dopamine responses in TS are correlated overall with approach behaviors, but not with either the current approach type or the next approach type on a trial-by-trial basis.

Taken together, our recording results reveal that dopamine release in TS correlates with approach types, with smaller responses correlating with individual engagement. However, the specific level of dopamine release in TS was not correlated with the current approach type, suggesting that acute dopamine concentration in TS does not directly regulate retreat behaviors, but rather sends a feedback or evaluation signal during risk assessment.

### Reinforcement learning model with a shaping bonus and uncertainty for novelty response

Our previous work suggested that dopamine in TS signals a threat prediction error (Menegas et al., 2018). However, it was not clear how dopamine signals algorithmically relate to novelty-driven behaviors. We therefore sought to develop a simple model that captures novelty-triggered behaviors in our data obtained for the 4 experimental groups (novel object, unexpected familiar object, sham and ablation) as well as dopamine release in TS. In standard reinforcement learning models, dopamine is typically modeled as temporal difference (TD) error. This is the difference of reward prediction (or value) of adjacent states, which is used as a teaching signal for incremental learning of reward predictions (Sutton and Barto, 2015). Using similar logic, we first modeled simple threat prediction learning with TD error (Figure 7). In this model, trials are denoted by bouts of approach towards and sampling of the object (Figure 7a). Because TS dopamine is activated upon encountering a novel stimulus, we added ’threat’ at the time when an agent reached the object (’object location’ hereafter; Figure 7b, far left), instead of adding reward as in reward prediction learning. Threat prediction is used to determine future behavioral choice by comparing prediction with a constant threat threshold. If the current threat prediction is lower than the threat threshold, an agent can engage. If threat prediction is higher than threat threshold, an agent should avoid (Figure 7b, far right).

**Figure 7.**
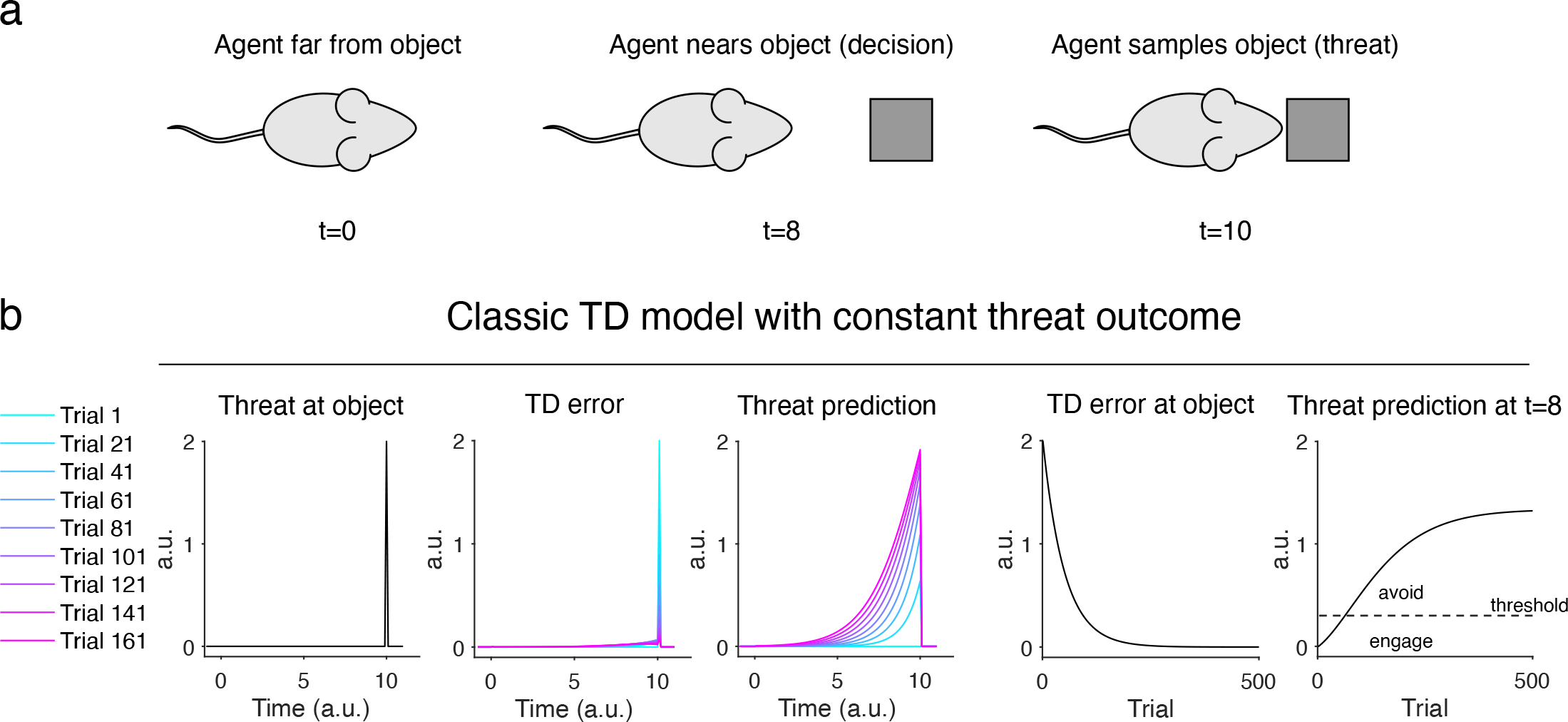
Basic reinforcement learning model with constant threat. **a.** Model approach-retreat bouts (equivalent to “trials” in traditional behavioral paradigms) are initiated independently by agent. As agent nears object, it makes a decision as to what behavior it will exhibit (risk assessment, engagement, or avoidance). An agent samples object features, which causes threat outcome. **b.** Temporal difference (TD) learning model that incorporates threat prediction (instead of reward prediction). Far left: constant threat signal is added at t=10 when agent reaches object. Second from left: Threat signal creates TD error at object location (t=10) in early trials. Middle: Threat prediction increases over trials as TD error decreases (threat becomes predicted). Second from right: TD error at object location decreases across trials as threat becomes predicted. Far right: Threat prediction propagates backward to earlier time points within bout such that threat prediction near object increases across trials. As agent nears object, it compares threat prediction to its own threat threshold (dotted line): if prediction remains below threshold, agent decides to engage with object. If prediction exceeds threshold, agent decides to avoid object from then on.

In this model, TD error shows a positive response at the object location, which gradually decreases over many encounters (Figure 7b, second panel from right). This is consistent with our observation that TS dopamine responses at the retreat onset gradually decreased (Figure 6e). The decrease of TD error is solely because threat is more predicted, thus generating a smaller prediction error, but the level of threat assigned to the object is kept constant (Figure 7b, far left). We then examined how threat predictions developed near the object location. Threat prediction before an agent reaches the object location gradually increased over multiple encounters and eventually plateaus (Figure 7b, far right). This tendency does not depend upon the amplitude of the threat (not shown). Because the threat threshold is a constant, the increase of threat prediction translates into a behavioral change from approach to avoidance (Figure 7b, far right). While increasing threat prediction explains the later avoidance exhibited by some animals in the novel object group, this explanation is inconsistent with our observation that some animals in the novel object group eventually showed engagement. Further, it does not explain why animals engage with familiar objects if the object is threatening.

We previously found that TS dopamine responses to a novel stimulus decayed when not associated with an outcome, whereas this decay slowed when it was associated with an outcome, especially a threatening outcome (Menegas et al., 2017). In this case, a novel stimulus can be interpreted as a threat-predicting cue instead of unconditioned threat stimulus. From these results, we proposed that threat prediction is initially high in TS dopamine (Menegas et al., 2017). We next expanded this idea to formally model dynamics of threat prediction in response to a novelty encounter using TD learning. We modeled threat learning with a positive default value of threat prediction assigned to a novel object, similar to a “shaping bonus” (Kakade and Dayan, 2002). A fixed value for the shaping bonus functions as a preliminary, initializing value of threat prediction, which speeds up (’shapes’) but does not distort eventual learning. In our model, an agent would eventually learn no outcome (no threat) associated with a novel object, but in the meantime, threat prediction and behaviors would be shaped by the initial estimation of threat prediction.

We examined the dynamics of TD errors and threat prediction using different levels of shaping bonus (i.e. initial threat prediction level) (Figure 8). The shaping bonus was applied at the object location to model a tentative guess of threat prediction according to the sampled sensory features without knowing the ultimate outcome. The actual outcome is eventually found out to be nothing (no threat) after the sampling behavior in each approach bout (Figure 8a). Because of the positive initializing value of threat prediction (Figure 8b, first column ’shaping bonus’), positive TD errors were initially observed at the object location, which then decreased over many encounters (Figure 8b, second and fifth columns), similar to the model with threat outcome (Figure 7).

**Figure 8.**
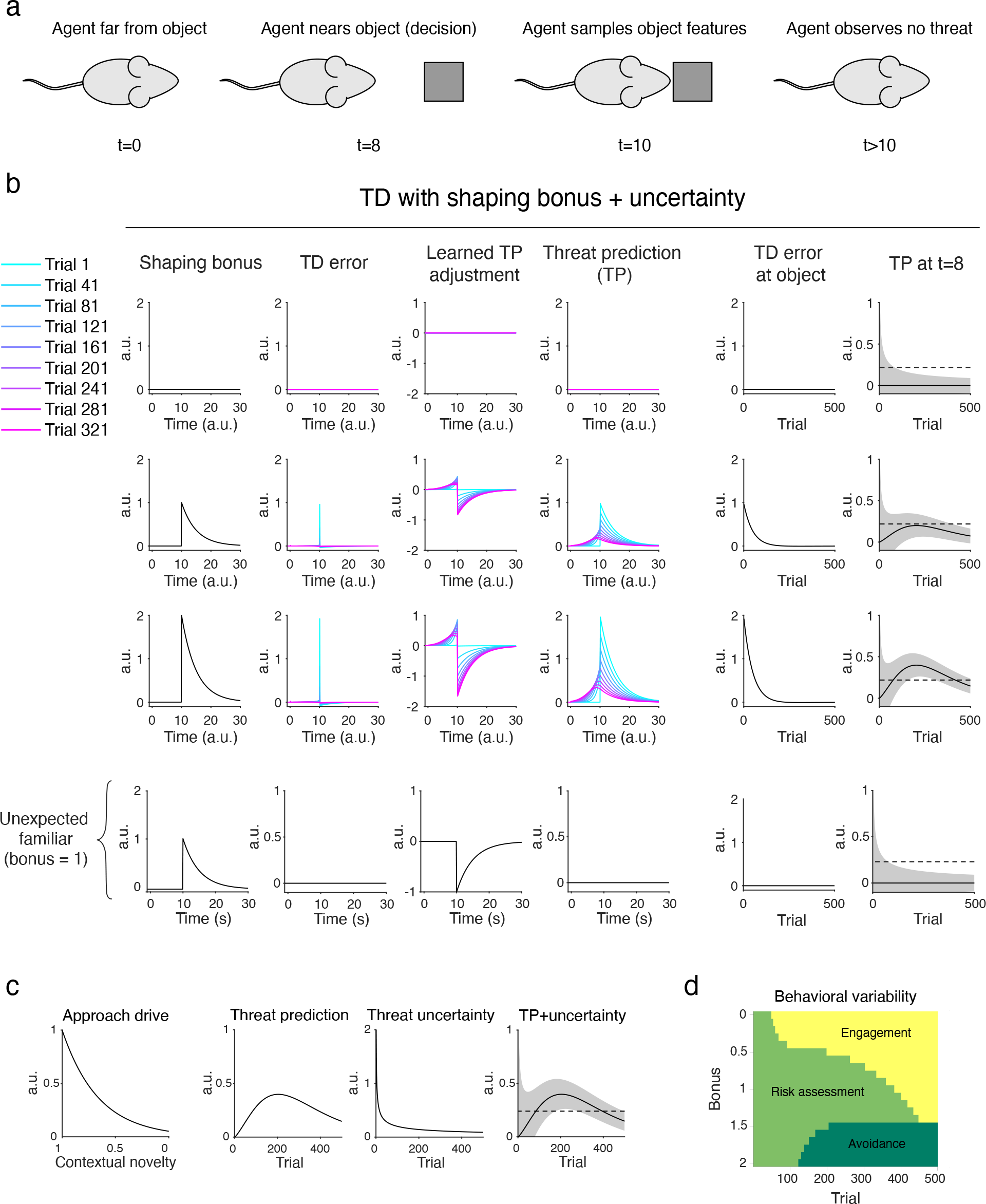
Reinforcement learning model with shaping bonus and uncertainty. **a.** Similar bout structure to Figure 7a, but agent never observes actual threat. **b.** TD learning model that incorporates both threat-shaping bonus and threat uncertainty. Instead of adding threat (Figure 7), model adds initial estimation of threat (far left column, ’shaping bonus’). The shaping bonus initially creates TD error at the object location (second column from left, cyan), which in turn reinforces threat prediction near the object (third column from left, t=8). The threat prediction (fourth column) is determined by summation of the initial estimation (shaping bonus, first column) and the learned component (third column). Threat prediction at object (t=10) starts out high, but decreases across trials since the sensory features, initially predicting threat, are not associated with actual threat (fourth column). TD error at object also decreases across trials (fifth column). Threat prediction near object (t=8) gradually increases at first, and then gradually decreases to 0 (fourth and far right column). Threat uncertainty is indicated by shading in the rightmost column. When an agent encounters unexpected familiar object, a shaping bonus is already canceled out by learning, but threat uncertainty causes initial risk assessment (bottom row). **c.** Components to determine behaviors. Left: Approach drive as a function of degree of contextual novelty (surprise). Approach drive potentiates approach, independent of threat prediction. Second from left: Threat prediction near object (t=8). Threat prediction determines whether to engage or avoid. Second from right: Threat uncertainty near object. Uncertainty of threat estimation was determined incrementally using a Kalman filter (see Methods). Threat uncertainty determines the degree of risk assessment. Right: Threat prediction plotted together with threat uncertainty, with uncertainty represented by shading. Dotted line representing threat threshold (same format as Figure 8b, rightmost column). **d.** Development of behavior based on different degrees of shaping bonus for novel object. Low shaping bonus (bonus=0) results in early transition from risk assessment to engagement, whereas high shaping bonus (bonus=2) results in transition to avoidance. Intermediate shaping bonus leads to prolonged risk assessment.

However, dynamics of threat prediction were different. Threat prediction at the object location was defined by the shaping bonus (Figure 8b, fourth column, cyan) and gradually decreased over trials to 0 (Figure 8b fourth column at time 10), because the actual outcome is nothing. In other words, the agent’s initial guess of threat prediction associated with the sensory features was wrong and subsequently updated by learning (Figure 8b, third column).

In the meantime, the threat prediction near the object increases initially because of positive TD errors caused by a shaping bonus, then decreases afterwards and eventually becomes 0 after learning has finished (Figure 8b, fourth and far right columns). Across different conditions as the shaping bonus increases, the peak of the threat prediction near the object increases, whereas the time-course is similar (Figure 8b, far right). The concave shape of threat prediction development near the object explains approach agents who eventually engage with a novel object (Figure 8b, second row), and avoidance agents who first approach but ultimately avoid (Figure 8b, third row, see below for termination of learning with avoidance). Differences in the level of shaping bonus can thus produce different patterns of behavior throughout learning (Figure 8d).

However, animals do not choose behaviors based solely on threat prediction level. Even if their estimate of potential threat is low, they should be still cautious if the estimation is uncertain.

Animals should engage when they are confident about their safety, while they should continue risk assessment when they are uncertain whether they really are safe. We therefore added uncertainty of threat prediction to the model as another determinant of behavior. To implement uncertainty in a principled way, we used a Kalman filter to incrementally determine estimation uncertainty (see Methods), and plotted this together with threat prediction (Figure 8b, c). In these example, threat prediction is plotted with a 95% confidence range.

We find that the uncertainty of threat prediction explains initial risk assessment in many situations. With low initial estimation of threat, uncertainty of threat prediction is high at the beginning, inducing risk assessment behaviors, but the uncertainty quickly decays and allows a fast switch to engagement (Figure 8b, first row). Similarly, an unexpected familiar object causes an initial risk assessment because of threat uncertainty, but does not induce avoidance because the initial estimation of threat prediction with the object features is already canceled out by learning during pre-exposure (Figure 8b, bottom row). On the other hand, with high initial estimation of threat, uncertainty is high at the beginning, and then threat prediction increases, causing longer risk assessment (Figure 8b, second row). If threat prediction gets bigger than a threshold, agent chooses to avoid. Once it avoids, it loses a chance to further learn threat prediction that would eventually become 0, which results in persistent avoidance (neophobia) (Figure 8b, third row). Thus, the degree of shaping bonus may determine whether an agent becomes neophobic or not. This is different from forced familiarization in many studies (Menegas et al., 2018; Morrens et al., 2020; Ogasawara et al., 2021) and in the pre-exposure in our experiments. In forced familiarization, animals cannot totally avoid an object and so continue to learn and update threat prediction. Thus, most animals will eventually not be afraid of the object, because the threat prediction becomes 0 (the object is not threatening), although these stressful conditions may engage different systems in extreme cases (Corey, 1978; Rebec et al., 1997). Finally, we consider “approach drive” (Figure 8c, left) to potentiate approach as a separate system, independent of the threat prediction system. Approach drive in this study may simply decay alongside contextual novelty.

The shaping bonus in this model is determined by the initial responses of dopamine in TS to an object, and initial responses vary by individual. Our previous studies found that responses of TS-projecting dopamine neurons are monotonically modulated with the physical salience (intensity) of an external stimulus in the environment (Menegas et al., 2018). Thus, representation of physical salience in TS dopamine will determine the shaping bonus in this model, which in turn facilitates development of threat prediction and affects future actions. Taken together, these results suggest that behavioral engagement with a novel object is well captured by a reinforcement learning model with a shaping bonus, one in which threat prediction builds up according to representation of physical salience of the object in TS dopamine. The combination of threat prediction and uncertainty determines alertness or cautiousness, which guides whether an agent should engage, avoid, or continue risk assessment. By changing the level of shaping bonus, which can be inferred from the level of TS dopamine, the model predicts the diverse and dynamical patterns of behaviors observed across individuals and experimental conditions.

## DISCUSSION

In this study, we propose a reinforcement learning model that captures behavioral dynamics and variability in response to novelty and incorporates dopamine in the tail of the striatum (TS). We were led to this model by examining novelty-induced behaviors in freely-moving animals using supervised (DeepLabCut) and unsupervised (MoSeq) machine learning tools. These approaches demonstrate that all mice exhibited risk assessment behaviors toward a novel object initially, followed by engagement or avoidance. These observations were corroborated by an independent analysis based on the animal’s postures using an unsupervised behavioral classification method (MoSeq). This found that syllables that are enriched at the beginning of a novel object exploration correspond to cautious approach and cautious retreat which together constitute a set of risk assessment behavior. Thus, our application of machine-learning based analysis methods allowed us to identify distinct behavioral motifs that are dynamically driven during an encounter with a novel object.

The observed distinct approach behaviors depart from the previous studies that categorized novelty-induced behaviors merely by two opposing choices (approach versus avoidance) along a single dimension. By distinguishing the approach types, we found that stimulus novelty and dopamine in TS specifically suppress post-assessment engagement, but not risk assessment. We used temporal difference (TD) learning to explain commonalities and differences in the behaviors from four experimental conditions: unexpected novel vs familiar object conditions, and ablation of TS-projecting dopamine neurons vs sham conditions. We constructed a simple model by incorporating an initializing value for threat prediction (shaping bonus), and uncertainty of threat prediction. In this model, TS dopamine, which conveys a threat prediction error, gradually builds up threat prediction over multiple encounters with a novel object. This in turn suppresses transition from the risk assessment phase to post-assessment engagement, causing neophobia in extreme cases. Thus, in contrast to classical animal behavior models of novelty, neophobia can be caused by development of threat prediction rather than novelty detection per se. As the object turns out not to be threatening, threat prediction gradually decreases which models habituation. In this way, the model captured not only the temporal dynamics of novelty responses, but also individual variability in the behaviors. Importantly, we found that variability in TS dopamine responses corresponded to individual variability in behavioral responses, providing a neural readout of shaping bonus for threat learning. Together, our findings provide insights into the computations and neural mechanisms that may underlie the dynamics of novelty-induced behaviors, including neophobia.

### Shaping bonus and neophobia

Novelty drives both immediate behavioral responses and learning. Novelty-triggered behavioral and neural responses have been intensively studied, and various computational models incorporate novelty components to understand optimal strategies and animal behaviors, because the generation of appropriate novelty responses has been linked to behavioral strategy and learning in daily life (Jaegle et al., 2019; Kakade and Dayan, 2002). While most computational models have focused on the approach aspect of novelty responses, our study has extended these ideas to model approach suppression by incorporating a shaping bonus and uncertainty into a reinforcement learning framework.

Learning an appropriate action is often difficult, because the action is too complicated to learn at once and because an action and its outcome are too temporally separated to easily establish causality. Therefore, in operant conditioning, it is often the case that behaviors are “shaped by making the contingencies of reinforcement increasingly more complex” (Skinner, 1975). In machine learning, some powerful learning models are often slow. To make learning more efficient and fast, some models have copied the idea of shaping from psychology by adding an extra reward (“shaping” or sometimes called an intrinsic reward) at an intermediate step for learning of longer sequential choices (Ng et al., 1999). However, adding an extra intermediate reward distorts the eventual learning; an agent might learn to acquire only the mid-point reward, preventing from learning from the actual reward in the future. To overcome the problem of learning distortion, a specific form of shaping (“potential-based shaping”) has been proposed (Ng et al., 1999). In this method, instead of adding an extra reward, reward expectation is added at an intermediate step to preserve original reward function, but still also “shapes” an agent’s actions and learning. In this way, potential-based shaping is equivalent to optimistic initialization of reward prediction (Wiewiora, 2003). As a consequence, reward prediction of a state is initialized with a positive value even before an agent has visited that state.

Optimal initialization of control systems plays a critical role not only in machine learning but also in animal behaviors. For example, animals can avoid some threatening stimuli using species-specific defensive systems even if they have never encountered them. On the other hand, animals can also learn not to avoid some stimuli that were previously threatening. These phenomena can be interpreted as an initialization of threat prediction with a pre-programmed value. In addition to these pre-programmed mechanisms, the initializing value could in principle be set by experiences in other states without visiting the actual state. Such flexible initialization is critical for efficient machine learning (“Smart initialization”, Simsek et al., 2011), and for behavioral choices in daily life, where agents/animals continuously face novel states. Rather than starting from uniform estimation over all states, an initial guess (generated via generalization) can help to quickly learn more accurate estimation.

In reinforcement learning, approach to novel objects or cues is often modeled using a “novelty bonus” or “shaping bonus”. We adapted this approach to model avoidance of a novel object. Our model differs from previous animal behavior models of novelty where fear is simply a decaying function with novelty (Blanchard et al., 1991; Gordon et al., 2014; Halliday, 1966; Hogan, 1965; Hughes, 1997; Lester, 1967; Montgomery, 1955; Thorpe, 1956) in that it predicts that threat prediction first builds up and then (potentially) decays. These dynamics explain a variety of observed behavioral patterns. We also incorporated uncertainty of the threat prediction into our model, thereby accommodating threat predictions ranging from risk assessment to engagement. While we used a frequency-based simple Kalman filter to compute uncertainty, other methods *−* such as those based on actual threat prediction distribution *−* could be applied (Dabney et al., 2020; Lowet et al., 2020). Interestingly, we found a unique phenomenon specific to threat learning. Once an agent learns that the object is threatening, an agent avoids the object entirely and loses a chance to further learn. As a consequence, the agent gets trapped in an avoidance state. Thus, our model changes the way we interpret neophobia. Neophobia may not be simply driven by abnormal novelty detection per se, but instead forms dynamically in two steps. Uncertainty of safety induces initial risk-assessment, which is followed by a learning process about which objects should be avoided.

Since neophobia was thought to be linked to novelty, brain areas engaged during neophobia have been proposed to be involved in novelty detection. In this study, we found that TS dopamine plays a role in neophobia. While we cannot exclude possibility that TS dopamine is involved in novelty detection, TS dopamine likely signals the physical salience (such as intensity) of external stimuli. Activity of dopamine in TS is initially correlated with the intensity of novel stimuli (Menegas et al., 2018) and then gradually decays depending on associated future events. If no event is associated with a novel stimulus, the response to the stimulus quickly decays by repeated representation, while if it is associated with an outcome, potentially with physical salience of the outcome, the response is sustained (Menegas et al., 2017). Thus dopamine responses in TS, instead of detecting novelty, are initialized depending on stimulus salience, and then responses are adjusted afterwards. Our model further predicts that TS dopamine excitation with positive initialization (’potential threat’) is used as an evaluation signal for learning of threat prediction at an earlier time point (before approach), which in turn prevents animals from approaching a potential threat. If TS dopamine responses to an object are high, threat prediction will be developed and animals will not approach anymore. In this way, TS dopamine system uses physical salience of a stimulus as a default value of threat prediction to shape defensive behaviors even before animals learn the exact threat level.

Consistent with this idea, ablating TS-projecting dopamine neurons caused premature engagement. Furthermore, we found that individual variability of dopamine activity in TS at least partially explains natural individual variability of level of neophobia. Notably, the individual variability of dopamine responses in TS may derive from variability in the initializing value, i.e. scoring of stimulus salience, rather than novelty detection. Hence, neophobia may be caused by abnormal threat prediction due to general sensitivity to sensory stimuli, rather than aberrant novelty detection.

Why, then, do animals avoid a novel salient stimulus in the first place? A recent series of studies found that in appetitive situations, the taste of food is not an ultimate outcome but instead functions as prediction of nutrients, which are the ultimate outcome or goal for eating (Fernandes et al., 2020; Han et al., 2018; Tellez et al., 2016). From these results, Dayan proposed that taste is a kind of shaping, an initial guess for value of eating, which can be updated according to an actual outcome, i.e. nutrients (Dayan, 2021). In this framework, dopamine responses to food rewards (taste) are tentative feedback based on shaping bonus, but not ultimate reward outcome, to facilitate learning. Similarly, some dopamine neurons are excited by novel odors but only when the animal preferred that odor behaviorally (Morrens et al., 2020), suggesting that odors may also function to initially estimate value. We can interpret our data in threat prediction with analogy to the idea in appetitive value (Figure S6). Our results suggest that there is another type of shaping that aids threat prediction without visiting the actual state. Similar to well-known pre-programed threats such as looming stimuli and predator odors, physical salience of stimuli may help animals to estimate threat without actual experiences. While many salient stimuli end up being non-threatening, caution against exploring high intensity novel stimuli may be lifesaving. Physical salience can be easily and quickly computed and easily generalized. Therefore, animals may routinely use physical salience as an initial guess of a potential threat for an immediate action and learning, because learning threat only from ultimate outcomes such as pain, injury and death may come at a high cost. Thus, the idea of shaping can be broadly applicable, and dopamine neurons with distinct activities can share a common framework.

### Diversity of dopamine neurons

While the role of dopamine in reward prediction has been relatively established (Eshel et al., 2013; Glimcher, 2011; Schultz, 2016; Watabe-Uchida and Uchida, 2018), our knowledge of functional diversity of dopamine neurons is still incomplete (Cox and Witten, 2019; Watabe-Uchida and Uchida, 2018). In particular, it is not yet clear whether non-canonical dopamine signals can be understood in the similar theoretical framework or algorithm as those in reinforcement learning theories. In our previous studies, we found that TS-projecting dopamine neurons do not signal rewards but respond to a set of threatening stimuli, including high intensity or novel stimuli (Menegas et al., 2017, 2018), and play a role in avoidance of them (Menegas et al., 2018).

Based on precise observation of behaviors and dopamine signals in response to novelty, we have obtained a clearer view on how TS dopamine functions during novelty exploration. First, it should be noted that, unlike previous experiments (Cohen et al., 2012; Menegas et al., 2017; Schultz et al., 1997; Tsutsui-Kimura et al., 2020), our work involves animals freely interacting with an environment. Nonetheless, discrete approach-retreat bouts in our novelty paradigms can be regarded as being equivalent to “trials” in more structured behavioral paradigms, albeit with a critical difference in that the animal can control “task” structure. Our results support the possibility that non-canonical dopamine signals found in TS work as an evaluation signal even in a naturalistic setting, in a manner similar to canonical dopamine signals observed in many structured tasks (Cohen et al., 2012; Glimcher, 2011; Schultz, 2015) or during social interactions (Dai et al., 2021; Gunaydin et al., 2014). Further, dopamine in TS, while signaling totally different information from canonical dopamine, may facilitate threat prediction in a similar manner that canonical dopamine facilitates reward prediction.

Together, our results suggest a possibility that even if information contents are diverse, the function of dopamine neurons can be understood within the common framework of reinforcement learning including an idea of bonuses for fine tuning.

## ACKNOWLEDGEMENTS

We thank Adam Lowet, Malcolm Campbell, and all lab members for discussion and feedback. This work was supported by the NIH BRAIN Initiative (U19NS113201, NU, SRD; and R01NS108740, NU), National Institute of Mental Health (R01MH125162, MW-U) Simons Collaboration on the Global Brain (NU, SRD), Bipolar Disorder Seed Grant Program (NU), Japan Society for the Promotion of Science (IT-K) and the Harvard Molecules, Cells, and Organisms Program training grant (KA). We thank the staff of the Harvard Center for Biological Imaging, Center for Brain Science Fabrication Lab, and the Physics/SEAS Machine Shop for technical support.

## AUTHOR CONTRIBUTIONS

KA, NU and MW-U initiated the project and designed the experiments. KA and YX set up the arena and wrote analysis code. AM and MWM assisted in DeepLabCut setup. JM, RA, and SRD assisted in MoSeq setup. KA trained the mice and collected the data. KA and IT-K performed surgery and histology. KA and MW-U analyzed the data and wrote the paper. KA, IT-K, YX, AM, MWM, NU, SRD and MW-U edited the paper.

## METHODS

### Animals

78 adult male and female mice were used. Behavioral experiments were performed on C57BL/6J mice (Jackson Laboratories), aged 9-17 weeks, on the dark cycle of a 12-hr dark/12-hr light cycle (dark from 7:00 to 19:00). Behavioral tests and recordings were conducted between 8:00 and 18:00. Animals were group-housed until testing or surgery, then individually housed throughout testing. All procedures were performed in accordance with the National Institutes of Health Guide for the Care and Use of Laboratory Animals and approved by the Harvard Animal Care and Use Committee.

### Behavioral apparatus

To assess naturalistic behaviors in mice, an open-field arena was developed that allowed the recording of free movement (see Table 1 for parts list). Mice were able to explore freely in a 60cm by 60cm flat arena, either empty or containing a single novel object in one corner. To record movement, a single camera was mounted on a beam ∼70cm above the floor of the arena. A bright white LED light (Westek Indoor Outdoor White LED Rope Light) illuminated arena from above.

**Table 1.**
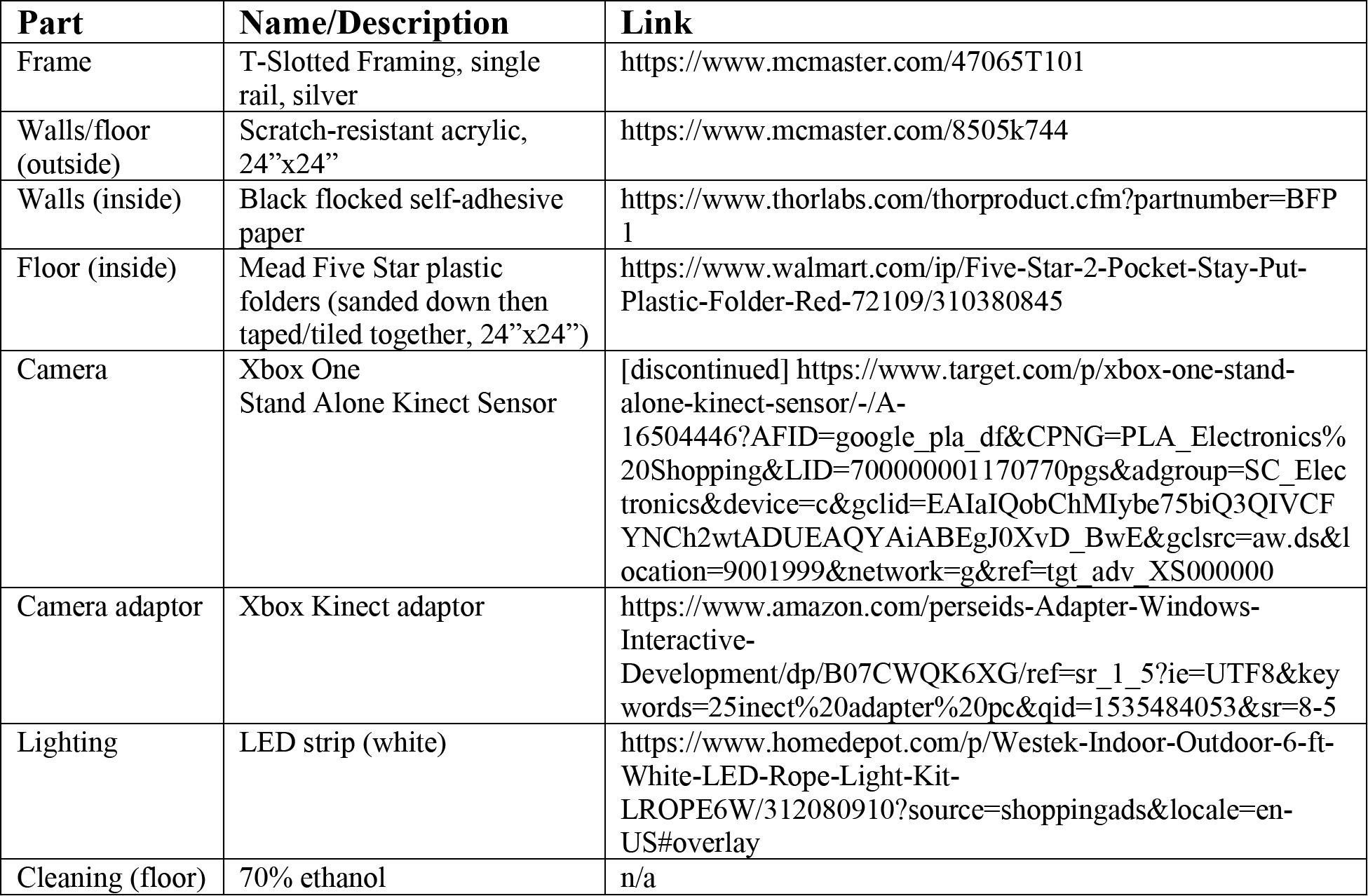
Parts list for novelty arena.

**Table 2.** Key resources table.

| Reagent type (species) or resource | Designation | Source or reference | Identifiers |
| --- | --- | --- | --- |
| Virus strain | Dopamine sensor | Vigene Biosciences (Sun et al., 2018, 2020) | AAV9-hSyn-GRABDA2m |
| Virus strain | tdTomato | UNC Vector Core | AAV5-CAG -tdTomato |
| Virus strain | GFP | UNC Vector Core | AAV5-CAG-GFP |

### Experiment workflow

Before start of experiment, mice were separated and individually housed at least 1 day in advance. Once separated, mice were then handled for 30 minutes per day for 3 days (see Handling). For a novel object/an unexpected familiar object tests, mice were then pre-exposed to the test object (or dummy object) in their home cage for 30 minutes per day for 4-7 days. Test objects were either legos (Mega Bloks First Builders 80-piece Classic Building Bag, 72 mice) or rubber dog toys (Kong Classic dog toy size M, 6 mice). Brand new test objects were used at the start of each set of mice (fresh out of packaging). Dummy objects were plastic coconut cups (Shindigz 16-oz Coconut Cups, 5.5-in tall). For each animal, the same object was used for the duration of the experiment (1 object per animal, each animal’s object was wiped with ethanol after every day). A novel object group and a sham surgery group were pooled for Figure 1.

### Handling

Handling consisted of weighing mice (on first day) and scooping mice into a transport box. This scooping was to acclimate a mouse to the way they would later be transferred from the behavioral arena back to their home cage. To scoop a mouse, the experimenter would hold a takeout box in a corner of the home cage, laying sideways with opening facing center of cage.

The experimenter would wait until the mouse approached and walked into the box before lifting the box up and tilting it gently upright. Then the box would be tilted back sideways, replaced onto the floor of the cage, and the mouse would be allowed to return to cage. If the mouse did not voluntarily approach the box within 10 minutes, the opening of the box would be moved closer to the mouse to encourage entry. Sessions lasted for 30 minutes or until the mouse was scooped at least 5 times, whichever occurred first.

### Pre-exposure

During pre-exposure, one object was placed in each mouse’s cage according to experimental condition (test object or dummy object). Each mouse’s object was kept consistent across pre-exposure days, with each object being wiped with ethanol between days. Sessions lasted 30 minutes per day for 7 days. During session, mouse would be allowed to explore object freely within home cage (including touching, moving, etc). Pre-exposure sessions occurred in dimly lit rooms.

### Habituation

During habituation, animals were placed in empty behavioral arena and allowed to explore freely. Mice were transferred from their home cage into the behavioral arena and transferred out of the arena by scooping with a takeout box. Behavior was recorded with a single overhead camera (Xbox Kinect; see Table 1 for materials list). Habituation sessions lasted 25 minutes per animal per day for 2 consecutive days. Mice were run in the same order each day (order determined randomly at the beginning of the experiment, and held constant for the rest of the experiment). If arena was soiled at the end of a session, feces would be removed and floor of arena would be spot cleaned using ethanol-soaked wipes before the next session began. Between rounds of experiments, arena was thoroughly cleaned and base of arena was wiped down with odorless eliminator (Ah! Products All Clear Odorless Odor Eliminator).

### Novelty testing

Novelty testing sessions consisted of animals exploring a single novel object within the behavioral arena. Object was placed in the corner of a behavioral arena (taped to floor to prevent animal from moving it, ∼12-15cm from either wall). Sessions lasted for 25 minutes per animal per day for 4-12 days, and mice were run in the same order as habituation each day. One object was used per animal for duration of experiment and the objects were not shared between animals. Before each session, object would be submerged in soiled bedding (mixture of bedding from each mouse’s cage in current round, 6 animals) and wiped off with dry kimwipe to remove excess bedding dust. Objects were wiped with ethanol after each day and allowed to air out overnight before use.

### Video recording and analysis

#### Recording

An Xbox One Kinect camera (Table 1) was mounted 70cm above the behavioral arena. Mice were videotaped with four channels: three color channels (RGB, 15fps) and one depth channel (30fps). The RGB video was used to locate body part locations (DeepLabCut) and the depth video was used to segment behavior (MoSeq). Data was saved using custom recording software (Wiltschko et al., 2015). Analysis code and instructions for running them are deposited on GitHub (https://github.com/ckakiti/Novelty_paper_2021).

### DeepLabCut analysis

For body part tracking, we used DeepLabCut version 1.0 (Mathis et al., 2018). Separate networks were used for different experimental settings: namely for mice without fiber implants (network A) and mice with fiber implants (network B). Both networks were run using a ResNet-50-based neural network (He et al., 2016; Insafutdinov et al., 2016) with default parameters for 1,030,000 training iterations. We provided manually labeled locations of four mouse body parts within video frames for training: nose, left ear base, right ear base, and tail base. For network A: We labeled 1760 frames taken from 64 videos. For network B: We labeled 540 frames taken from 17 videos. For both networks, 95% of labeled frames were then used for training.

After running DeepLabCut on each video file, we processed the output files (csv array with x/y coordinates and likelihood values for each body part). First, we trimmed the early frames that had low (<10%) likelihood values, indicating that the mouse was not present in the arena yet, or they had poor tracking. We then corrected “jumps” in tracking, defined as a >15cm/frame change in Euclidian distance. Points identified as jumps were replaced by the mean of the previous frame and the following frame. Jumps were corrected separately for each body part (nose, left ear, right ear, and tail). Trajectories for each body part were then smoothed using a lowess moving average filter (5 points, default).

A body part was determined to be “near” the object if it fell within a radius of 7cm (Euclidean distance) from the center of the object. An approach bout is defined as either the nose or tail entering near the object, and the end of this bout is determined when the nose and tail are no longer near the object. Habituation sessions did not have an object present; therefore the area of analysis was chosen based on the position where the object would be in later sessions. This radius was chosen to not be too large and include edge walking (since the object was placed near the corner) but also not to be too small and fail to capture enough of the animals’ trajectory.

These approach bouts can be further broken down into whether the nose was closer to the object than the tail for the entire bout (approach with tail behind) or whether the tail was closer at some point (approach with tail exposure). Frequency of tail behind or tail exposure were calculated based on the number of bouts with tail behind approach versus tail exposed approach. Retreat timing was determined to be the closest point of the nose relative to the object before the mouse moves away.

### MoSeq analysis

Raw imaging data was collected from the depth camera, pre-processed (filtered, background subtracted, and parallax corrected), and submitted to a machine learning algorithm that evaluates the pose dynamics over time (Wiltschko et al., 2015). During video extraction (moseq2-extract), 900 frames were trimmed from the beginning of the video to correct for time between when video was started and when the mouse was placed in arena. During model learning (moseq2-model), a hyperparameter was set to the total number of frames in the training set (kappa=2,711,134, 52 sessions, 52 animals). This exceeds the recommended >=1 million frames (at 30 frames per second) needed to ensure quality MoSeq modeling.

To align syllables to retreat timing, MoSeq data was aligned to DeepLabCut timeframes. This alignment was necessary because the depth and rgb videos have different frame rates (depth=30fps, rgb=15fps; timestamps are saved alongside raw data). We first extracted the timestamps and syllables associated with each frame in the depth video (scalars_to_dataframe function; see GitHub repository “moseq2-app”, code available on request: datta.hms.harvard.edu/research/behavioral-analysis/). We then aligned the depth video timestamps to the corresponding rgb video timestamps (custom MATLAB script, see GitHub repository “Novelty_paper_2021”). This alignment was then used to determine which syllables were expressed at each frame in the RGB videos. We then identified retreat timing and the corresponding MoSeq syllable in each RGB video.

### Surgical procedures

All surgeries were performed under aseptic conditions with animals anesthetized with isofluorane (1-2% at 0.5-1.0 l/min). Analgesia was administered pre-(buprenorphine, 0.1mg/kg, I.P.) and post-operatively (ketoprofen, 5 mg/kg, I.P.). At the time of surgery, mice were 2-4 months old. We used the following coordinates to target injections and implants for tail of striatum (TS): Bregma: -1.5 mm, Lateral: +3.0 mm, Depth: -2.4 mm (relative to dura) (Paxinos and Franklin, 2019)

### 6-OHDA surgical procedure

To bilaterally ablate dopamine neurons projecting to TS, we followed an existing protocol (Menegas et al., 2018; Thiele et al., 2012). The following solution was injected (I.P.) to animals at 10 mg/kg:

- 28.5 mg desipramine (Sigma-Aldrich, D3900-1G)
- 6.2 mg pargyline (Sigma-Aldrich, P8013-500MG)
- 10 mL water
- NaOH to pH 7.4

Most animals (weighing ∼25g) received ∼250 µL of this solution. This was given to prevent dopamine uptake in noradrenaline neurons and to increase the selectivity of uptake by dopamine neurons. After injection, mice were anesthetized as described above. We then prepared a solution of 10 mg/mL 6-hydroxydopamine (6-OHDA; Sigma-Aldrich, H116-5MG) and 0.2% ascorbic acid in saline (0.9% NaCL; Sigma-Aldrich, PHR1008-2G). The ascorbic acid in this solution helps prevent 6-OHDA from breaking down. Control animals were injected with vehicle ascorbic acid solution. To further prevent 6-OHDA from breaking down, we kept the solution on ice, wrapped in aluminum foil, and it was used within three hours of preparation. If the solution turned brown in this time (indicating that 6-OHDA has broken down), it was discarded and fresh solution was made. 6-OHDA (or vehicle, ascorbic acid solution) was injected bilaterally into TS (200nL per side). Each injection was spread out over several minutes (70-100 nl per minute) to minimize damage to the tissue. Surgeries occurred 1 week before handling.

### Dopamine sensor surgical procedure

For TS neurons to express dopamine sensor for fluorometry, we unilaterally injected mixed virus solution (AAV for dopamine sensor and tdTomato, 1:1 mixture, 350 nl total) into TS in WT mice. Virus injection lasted around 5 minutes (injection of 70-100 nl per minute), after which the pipette was slowly removed to prevent damage to the tissue. We also implanted optic fibers (400 µm diameter, Doric Lenses, Canada) unilaterally into the TS (one fiber per mouse). Once fibers were lowered, we attached them to the skull with UV-curing epoxy (Thorlabs, NOA81), then waited for 15 min for this to dry. We then added a layer of black Ortho-Jet dental adhesive (Ortho-Jet, Lang Dental, IL). We used magnetic fiber cannulas (Doric Lenses, MFC_400/430) to allow for recording in freely moving animals. We waited for 15 min for the dental adhesive to dry, and then the surgery was complete.

### Histology and immunohistochemistry

Histology was conducted in the same manner as previously reported (Tsutsui-Kimura et al., 2020). Mice were perfused using 4% paraformaldehyde, then brains were sliced into 100µm thick coronal sections using a vibratome (Leica) and stored in PBS. These slices were then stained with rabbit anti-tyrosine hydroxylase (TH; AB152, EMD Millipore) at 4°C for 2d to reveal dopamine axons in the striatum, dopamine cell bodies in the midbrain, and other neurons expressing TH throughout the brain. Slices were then stained with fluorescent secondary antibodies at 4°C for 1d. Slices were then mounted in anti-fade solution (VECTASHIELD anti-fade mounting medium, H-1200, Vector Laboratories, CA) and imaged using Zeiss Axio Scan Z1 slide scanner fluorescence microscope (Zeiss, Germany).

### Fluorometry (photometry) recording

#### Overview

Fiber fluorometry signal was recorded from the striatum in mice performing open field novelty behavior tasks (15 animals). Mice were injected either with AAV to express dopamine sensor. After undergoing surgery (details in Surgical Procedures), animals were allowed to recover for 2 weeks before the start of behavior testing. In the last 3 days of this period, animals were handled (details in Handling). Then animals went through habituation and novelty testing in the arena (described in a previous section). During photometry recordings, a long flexible optic fiber (see Recording section) was attached to connector on the animal’s skull which did not impede animal movement.

#### Handling

In addition to weighing and scooping mice in the takeout box, photometry mice also had a patch cord attached and removed once during the session (not connected to laser, no light transmitted). Animal was allowed to briefly move about cage with patch cord attached (∼10s) before being picked back up and disconnected from patch cord. Attachment and removal were conducted in same manner that they would be later in behavioral sessions.

#### Recording

Fluorometry recording was performed as previously reported (Menegas et al., 2018; Tsutsui-Kimura et al., 2020). The following describes this established setup: We use an optic fiber to stably access deep brain regions and interfaces with a flexible patch cord (3 m, Doric Lenses, Canada) on the skull. The patch cord simultaneously delivers excitation light (473 nm, Laserglow Technologies, Canada; 561 nm, Opto Engine LLC, UT) and collect dopamine sensor and tdTomato fluorescence emissions. Activity-dependent fluorescence emitted by cells in the vicinity of the implanted fiber’s tip (NA=0.48) was spectrally separated from the excitation light using a dichroic, passed through a single band filter, and focused on a photodetector connected to a current preamplifier (SR570, Stanford Research Systems, CA).

During photometry recording, optic fibers on the animal’s skull were connected to a magnetic patch cable (Doric Lenses, MFP_400/430) which both delivered excitation light (473 and 561 nm) and collected emitted light. The emitted light was then filtered using a 493/574 nm beam-splitter (Semrock, NY), followed by a 500 ± 20 nm (Chroma, VT) and 661 ± 20 nm (Semrock, NY) bandpass filters and collected by a photodetector (FDS10 X 10 silicone photodiode, Thorlabs, NJ) which is connected to a current preamplifier (SR570, Stanford Research Systems, CA). This preamplifier outputs a voltage signal which was collected by a NIDAQ board (National Instruments, TX) and custom Labview software (National Instruments, TX).

Lasers were turned at least 30 minutes prior to recording to allow them to stabilize. Before each recording session, laser power and amplifier settings were individually adjusted for each mouse. First, the laser power was set low enough to avoid bleaching and high enough to detect signal. Then, the amplifiers were set such that the baseline signals recorded through LabView were similar across mice and days (3-6 a.u. at start of session). Behavior and photometry signal were measured simultaneously using Labview software (see Synchronization section below). After each recording session, collected light intensity was measured from the patch cord using a photometer. Light intensity fell within a range of 15-180µW across animals and days.

### Signal analysis

DA sensor (green) and tdTomato (red) signals were collected as voltage measurements from current pre-amplifiers (SR570, Stanford Research Systems, CA). Green and red signals were cleaned by removing 60 Hz noise with bandstop FIR filter 58-62 Hz and smoothing with a moving average of signals in 50 ms. The global change within a session was normalized using a moving median of 100 s. Then, the correlation between green and red signals was examined by linear regression. If the correlation was significant (p<0.05), the fitted red signals were subtracted from green signals. Responses aligned at a behavioral event were calculated by subtracting the average baseline activity (-3s to -1s before the event) from the average activity of the target window (0-1s after the event).

### Synchronization

In order to match photometry signal to behavior, it was important to synchronize the rgb video and photometry data. To achieve this, an LED was mounted within view of rgb camera such that it appeared in video, but did not overlap the floor of the arena or obscure the mouse. Custom

LabView software was programmed to send a short TTL signal for a brief LED pulse every 10s for the duration of recording. TTL pulses and photometry signal were recorded simultaneously. After recording, the timing of LED flashing in the rgb video was determined and matched with the corresponding TTL pulses that had been saved alongside photometry signal. The result is two arrays of the same length: one containing the RGB frame number for each LED flash and the other containing the photometry timestamp for each TTL pulse (i.e. every 10s). The time for other frames were determined by evenly spacing those frames within 10s intervals.

### Statistical analyses

Data analysis was performed using custom software written in MATLAB (MathWorks, Natick, MA, USA). All error bars in the figures are SEM. In boxplots, the edges of the boxes are the 25^th^ and 75^th^ percentiles, and the whiskers extend to the most extreme data points not considered outliers.

### Modeling

#### Reinforcement learning of threat prediction

We applied the standard formulation of temporal difference (TD) learning (Schultz et al., 1997; Sutton and Barto, 1990) to threat prediction. In standard TD learning models (Sutton and Barto, 1998), an agent predicts the cumulative future rewards, or value. In our TD model, an agent predicts the cumulative future threats (threatening outcomes) to guide its behavior. We note that TD learning algorithm was originally developed for explaining the strength of association in a type of aversive conditioning (nictitating membrane response) (Sutton and Barto, 1987, 1990). There have also been some efforts to generalize TD learning algorithms to predictions of other quantity or outcomes (or “cumulants”) than value (Dayan, 1993; Schlegel et al., 2021). Our application of TD learning to threat prediction takes a similar approach to these precedents.

The threat prediction at time *t* is denoted as *TP*(*t*), and is defined by,

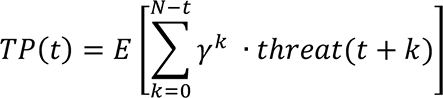

where *E*[… ] denotes expectation, *threat*(*t*) denotes a threatening outcome occurring at time *t*, and *γ* ∈ (0,1) is a discount factor. The model contained N (N=350) discrete states or timesteps, which constitute an entire bout of novel object exploration, with a novel object occurring upon entering to the 100^th^ state (*t* = 10) (for convenience, we express time *t* as the number of timesteps divided by 10). For simplicity, we applied a form of state representation called a complete serial compound, in which an agent deterministically traverses each of the 35 states in sequence (Schultz et al., 1997; Sutton and Barto, 1990), without considering avoidance action that would terminate state transitions and, thus, learning (see below).

In the first model (Figure 7b), we assumed that a threatening outcome occurred when the animal encountered a novel object (i.e. *t* = 10). That is, the novel object itself is a threat. Thus,

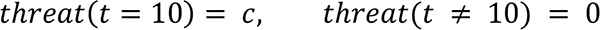

where *c* is a constant (in the Figure, *c* = 2 was used). Threat prediction, TP, was initialized to 0 for all the states before trial 1.

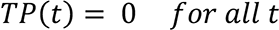

In each trial, the eligibility trace, *e*_*t*_, was initialized to 0 at the beginning of a trial. At each time *t*, TD errors, δ, were computed similar to a standard definition of TD error (Sutton and Barto, 1987) as the difference between the threat prediction at consecutive time steps plus received threats at each time step.

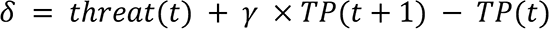

Eligibility trace, *e*_*t*_, for each state was updated by decaying *e*_*t*_ by the discount factor (*γ*) and the eligibility trace parameter (*λ*). For the current state, 1 was added.

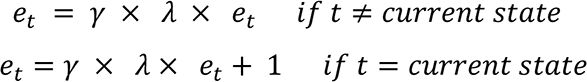

Threat prediction was updated according to the obtained *γ* and *e*_*t*_,

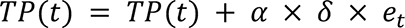

where α ∈ (0,1) is a learning rate. Then, an agent moves to the next time step, starting the next iteration of threat prediction. In this model, TD error at object (*t* = 10) is expressed as:

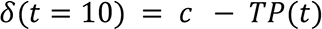

which is simply threat minus learned threat prediction.

The second model (Figure 7c) does not experience an actual threatening outcome but an initializing value (0 to 2) of threat prediction (i.e., shaping bonus Φ) was added to the state containing a novel object (*t* = 10) that gradually decays, to simulate lingering threat prediction until the animal finds out that there is no threat outcome. Thus, before starting the trial 1,

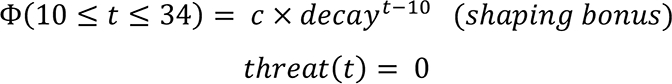

We used constant *c* from 0 to 2, and decay=0.98 in the Figure 8 Different levels of *c* yielded different time-course of threat prediction and prediction error in this model. Since Φ is an initializing value of threat prediction, threat prediction can be expressed as:

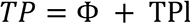

where TPl denotes learned component of threat prediction. Iteration of threat prediction was performed similarly to the model 1.

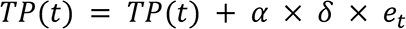

Since shaping bonus is fixed across trials, the learning rule can be also expressed as:

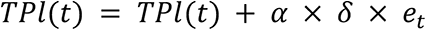

In this model, TD error is expressed as:

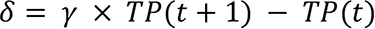

because there is no actual threat in any time step.

In all simulations, the learning rate *α*, the discounting rate *γ* and the parameter for eligibility trace *λ* were fixed to 0.02, 0.98, and 0.9, respectively, without model exploration.

### Uncertainty

Uncertainty of threat prediction (estimation uncertainty), *pp*(*n*), in each trial *n* was determined incrementally using the following equation (Kalman filter):

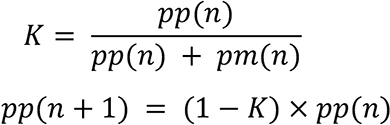

where *pm* is a measurement uncertainty. The model used standard normal distribution for estimation (threat prediction) in trial 1, and measurement (actual threat) in all trials, so that both variance *pp*(1) and *pm*(*n*) was set to 1.

### Behavioral choice

Behavior (risk assessment, engagement and avoidance) was chosen every time the agent entered the state near the object (*t* = 8), according to the threat prediction near the object, *TP*(*t* = 8) and uncertainty, *pp*(*n*), compared to a threat threshold, *thresh*.

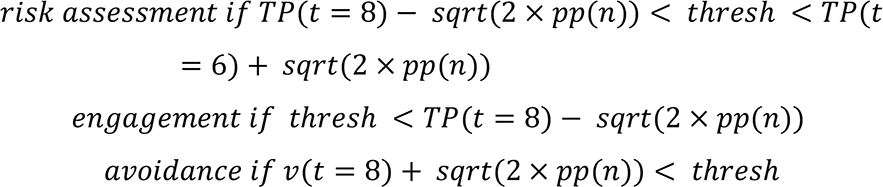

where engagement was chosen only if threat prediction is below threat threshold with > 95% confidence level. *thresh* = 0.2 was used for Figure 7e.

### Approach drive

In parallel with threat prediction and threat uncertainty, we showed approach drive as exponential decay function with decrease of contextual novelty.

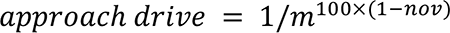

where *m* (*m*>0) is a constant and *nov* ∈ (0,1) is contextual novelty. We used m=1.03 for the Figure. Approach drive is independent of threat system, and we did not incorporate it in the present model.

### Data Availability

Matlab code files will be available on GitHub (https://github.com/ckakiti/Novelty_paper_2021). Video tracking and dopamine fluorometry data will be deposited.

---

**Figure S1.**
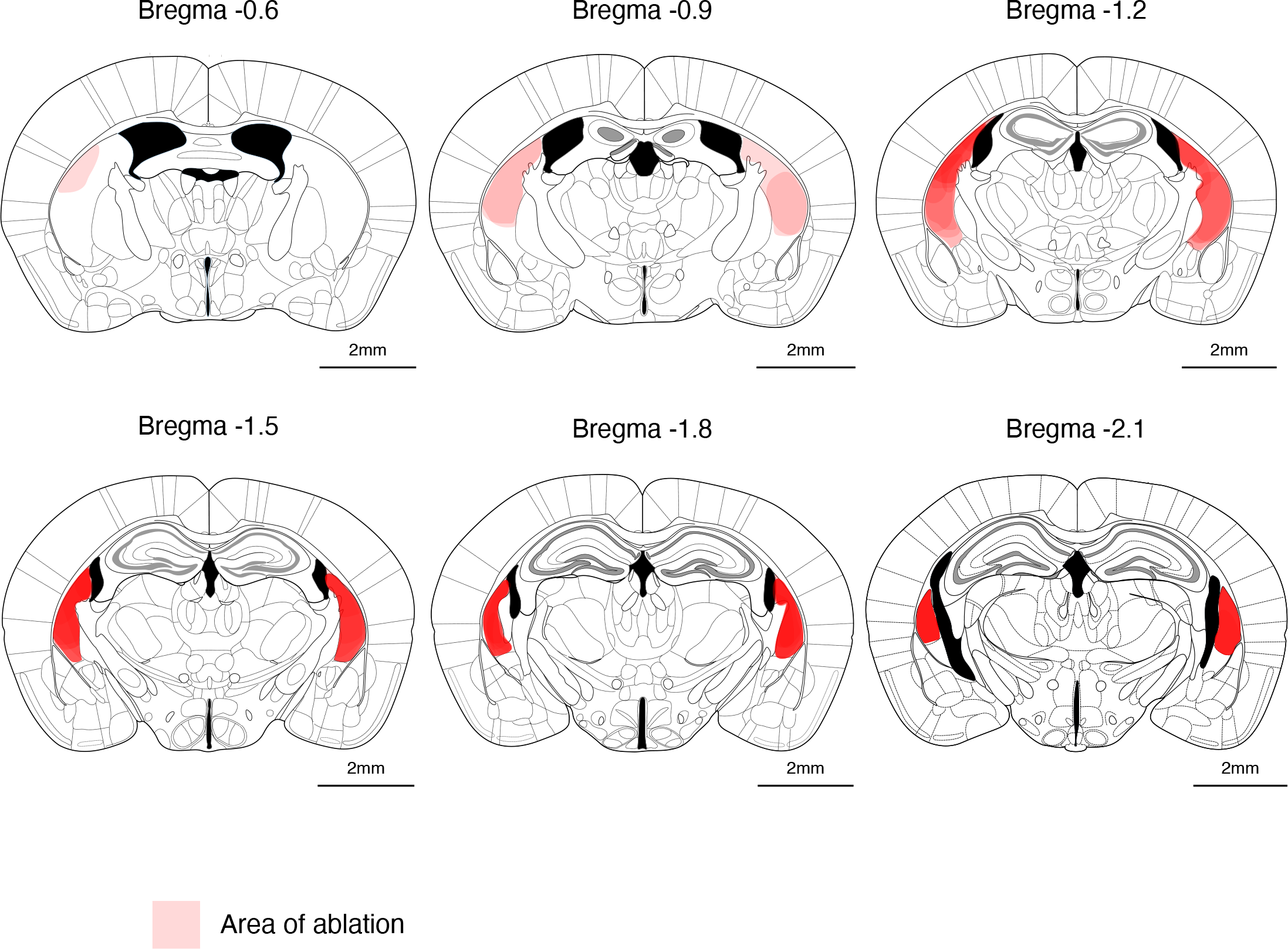
Summary of extent of TS-dopamine axon ablations. Extent of bilateral ablations of dopamine axons with 6OHDA (Figure 4) examined by immunohistochemistry with anti-tyrosine hydroxylase (TH) antibody. Ablation areas are marked with red shading, with overlapping shading from ablation mice (n=17). More densely overlapping areas have redder shading. Reference slices are depicted with 0.3 mm spacing (Paxinos and Franklin, 2019).

**Figure S2.**
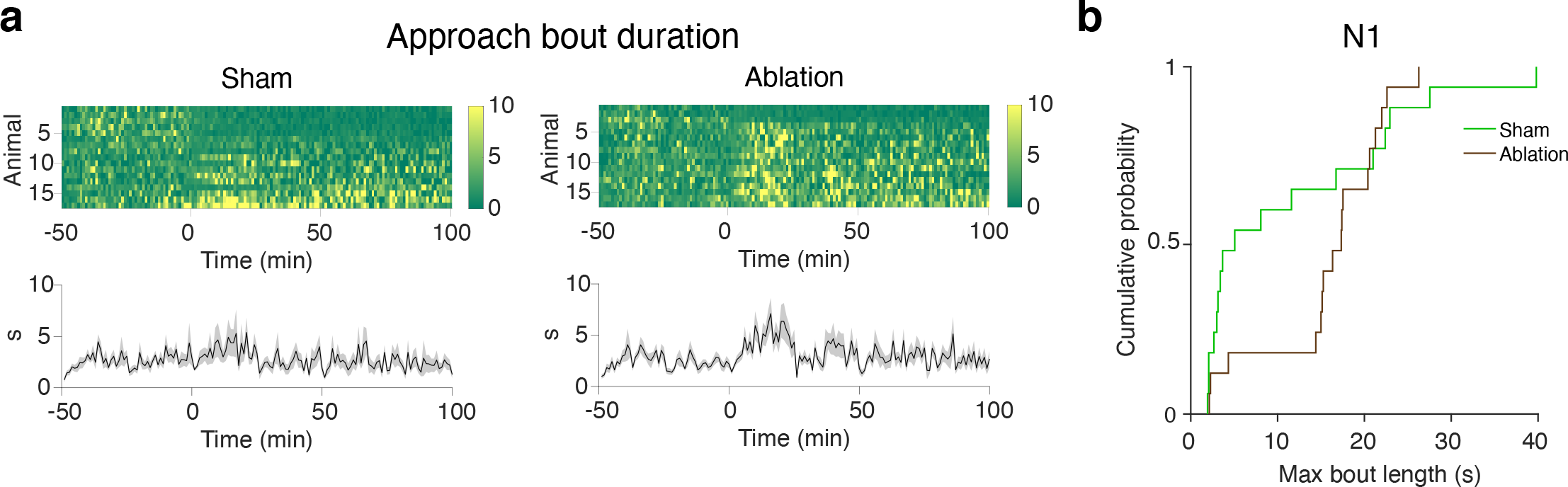
Ablation of TS-projecting dopamine neurons causes an increase in approach bout duration. **a.** Top: Duration of approach bouts in sham mice (left) or TS dopamine neurons ablation mice (right) (Figure 4). Ablation mice tend to exhibit longer bouts, especially on the first day of novelty. **b.** Cumulative probability of each group exhibiting different maximum bout lengths. Dopamine ablation causes maximum bout lengths to increase (p=0.030, n=17 for sham, n=17 for ablation, Kolmogorov-Smirnov [K-S] test).

**Figure S3.**
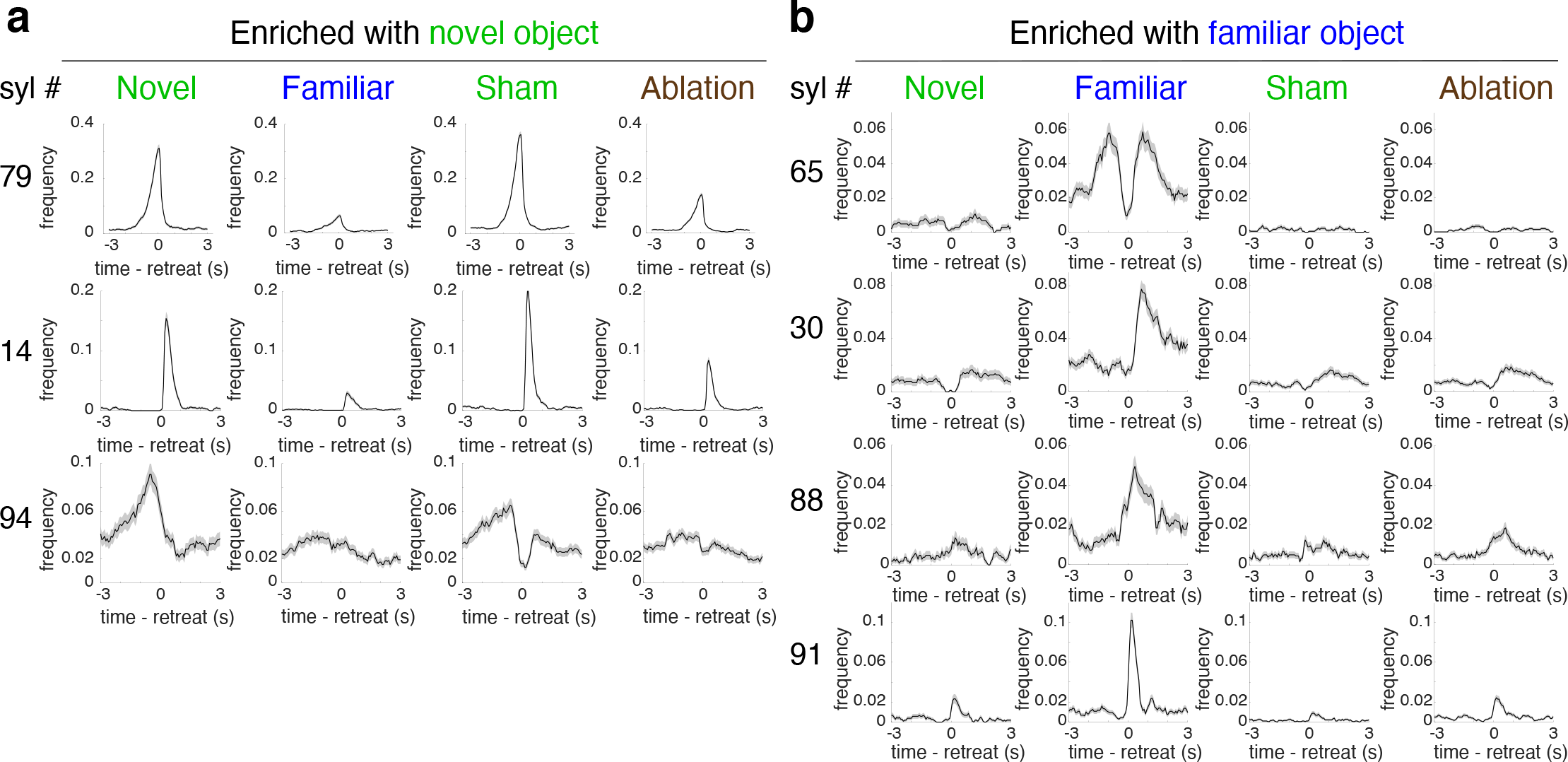
Distinct behavioral patterns in different experimental conditions were revealed with MoSeq. **a.** Frequency of retreat-aligned syllable usage for syllables that were significantly enriched with novel object (mean *±* SEM) (Figure 5). **b.** Frequency of retreat-aligned syllable usage for syllables that were significantly enriched with unexpected familiar object (mean *±* SEM).

**Figure S4.**
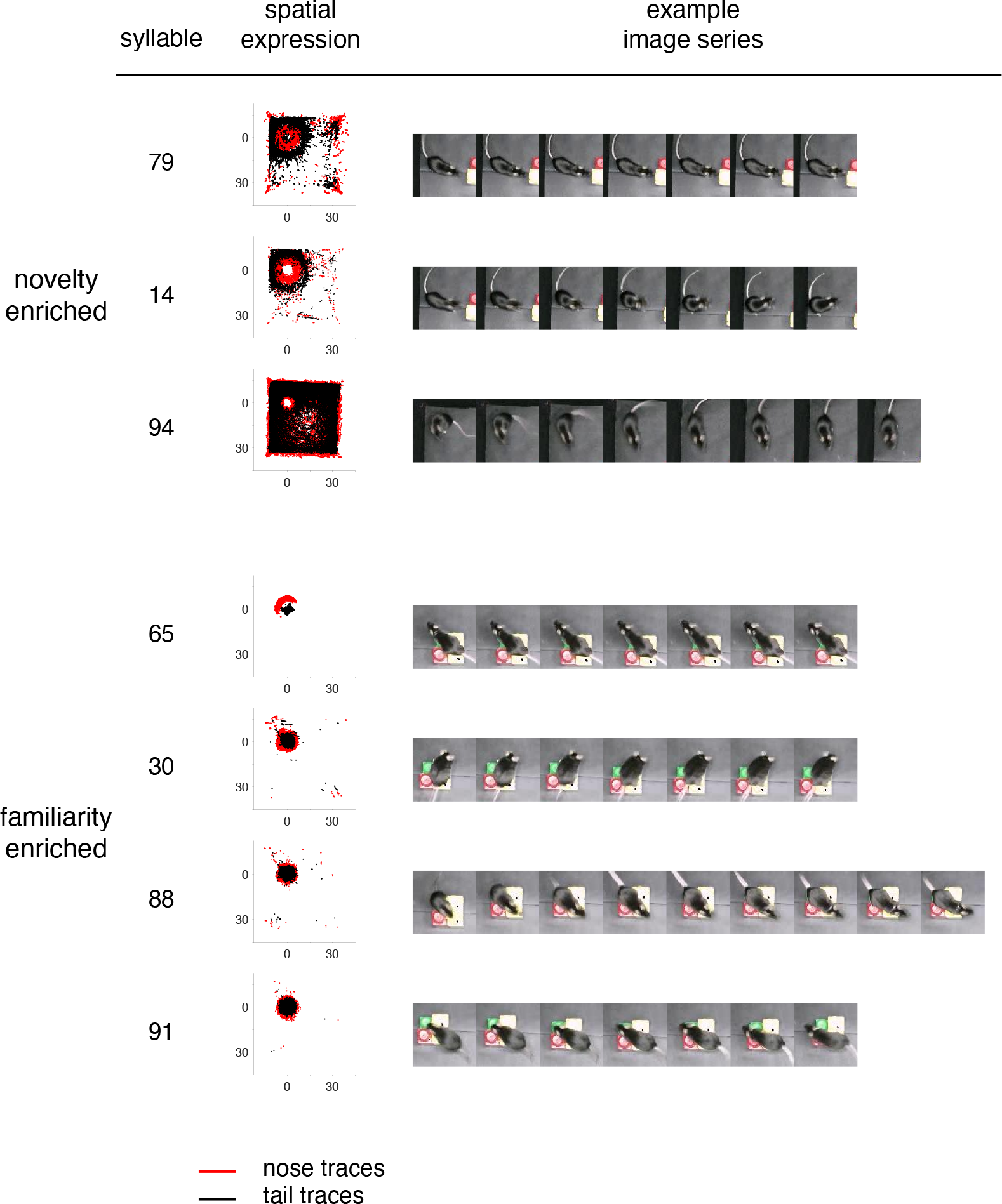
Characteristic postures enriched with a novel or unexpected familiar object. Left column: Syllables enriched in a novel object group and an unexpected familiar object group (Figure 5). Middle column: Spatial expression of each syllable. Nose trajectories are plotted in red and tail trajectories are plotted in black. Axes centered on object in top left corner (cm). Right column: Representative image series showing posture evolution of each syllable.

**Figure S5.**
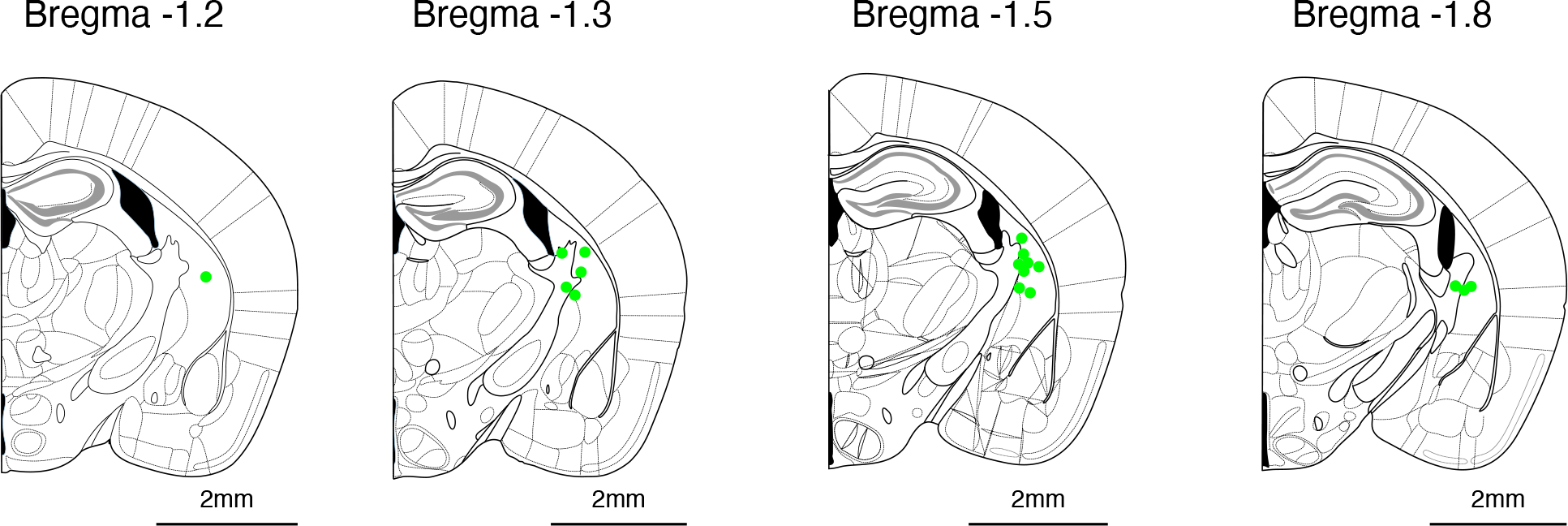
Summary of photometry recording fiber implant sites. Location of optic fiber tips used to collect dopamine sensor signals in TS (Figure 6). Fiber tip positions are plotted on the nearest reference slice (Paxinos and Franklin, 2019). Light emitted by neurons directly below these sites could be detected with photometry.

**Figure S6.**
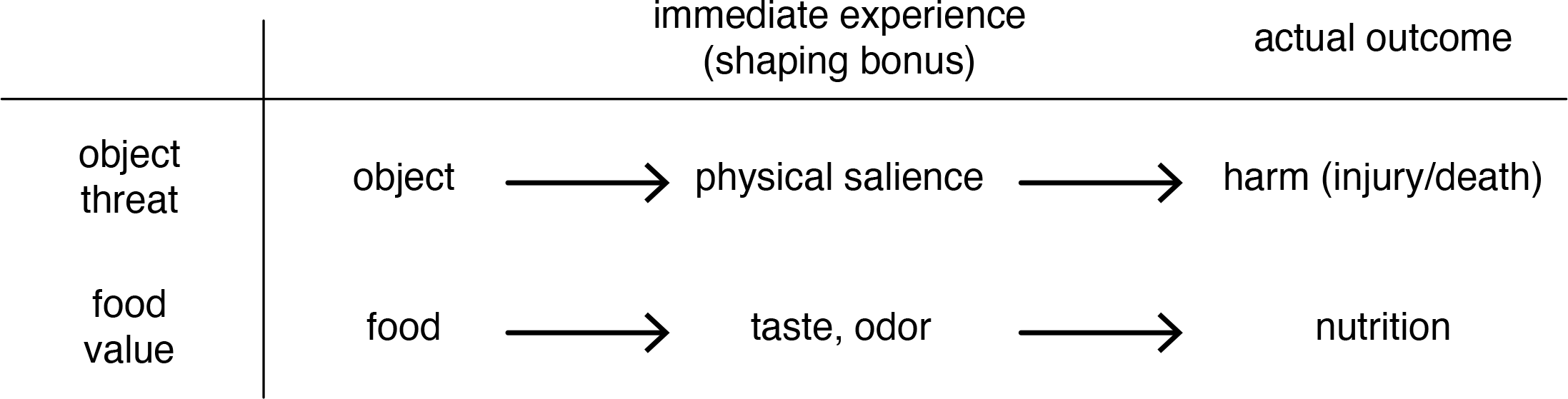
Use of immediate experiences as a preliminary estimation of the actual outcome. Comparison of sequence of events following encounters with food or object. Top row: Sequence of events for threat prediction. Animals may use physical salience to estimate actual harm in the future. Bottom row: Sequence of events for food value prediction. Animals may use taste or odor to estimate nutritional benefit or detriment. Shaping bonus models the estimation at time of immediate experience, before an animal has experienced the actual outcome.

Video S1 | Crowd video showing instances of syllable 79 expression.

Video showing 3D depth recording of behavior. Hotter colors indicate pixels that are closer to the camera. Object is in top left corner of each image. Multiple different instances of mice expressing the syllable are superimposed. A subset of instances (20) was randomly selected from the total instances for the given syllable, each with slight variation in duration. Video is time-locked such that the different syllable instances are expressed at approximately the same time during the crowd video. Mice are in light blue, and red dots indicate when mouse expresses the syllable of interest. Object is in top left corner of each image. A variety of instances of each syllable are shown, which together demonstrate the stereotypy within each syllable.

Video S2 | Crowd video showing instances of expression of syllable 14.

Same format as in Supplemental Video S1, shows syllable 14 instead.

